# Small RNA regulation of an essential process induces bacterial resistance to aminoglycosides during oxidative stress

**DOI:** 10.1101/2023.10.13.562219

**Authors:** Corentin Baussier, Charlotte Oriol, Sylvain Durand, Beatrice Py, Pierre Mandin

## Abstract

Fe-S clusters are essential cofactors involved in many reactions across all domains of life. In *Escherichia coli* and other enterobacteria, Fe-S cluster synthesis involves two machineries: Isc and Suf. While Isc functions as a housekeeping system, Suf is activated under stress conditions such as iron starvation or oxidative stress. Interestingly, cells functioning under Suf show reduced entry of aminoglycosides, leading to resistance to these antibiotics. The transcriptional regulator IscR, itself an Fe-S cluster containing protein, controls the transition between Isc and Suf machineries. Noteworthy, IscR has a critical impact on the virulence of various bacterial pathogens by regulating both Fe-S biogenesis and other pathways directly linked to host adaptation. Here, we discovered that two small regulatory RNAs (sRNAs), FnrS and OxyS, control *iscR* expression by base-pairing to the 5’ UTR of the *iscR* mRNA. Remarkably, these sRNAs act in opposite ways and in opposite conditions: FnrS, expressed in anaerobiosis, represses the expression of *iscR* while OxyS, expressed during oxidative stress, activates it. Using an *E. coli* strain experiencing protracted oxidative stress, we further demonstrate that *iscR* expression is rapidly and significantly enhanced in the presence of OxyS. Strikingly, we further show that OxyS induces resistance to aminoglycosides during oxidative stress through this unexpected regulation of Fe-S clusters biogenesis, revealing a new role for this sRNA.

**Significance Statement:** This study sheds light on the regulatory mechanisms controlling the synthesis of essential Fe-S clusters in bacteria, revealing unexpected roles for two small RNAs (FnrS and OxyS) in modulating the expression of the transcriptional regulator IscR. The findings suggest that this regulatory network could lead bacterial resistance to aminoglycoside antibiotics during oxidative stress, a condition associated with chronic infections. Ultimately, this work highlights the importance of understanding the intricate regulatory networks controlling bacterial metabolism and adaptation to stress, which could have significant implications for public health.

## Introduction

Adapting to ever-changing environments is a key requirement for all living organisms. Bacteria, in particular, have developed a remarkable ability to modify their physiology through genetic regulations, enabling them to quickly return to a state of balance. In recent years, small regulatory RNAs (sRNAs) have emerged as key actors in such adaptative responses in bacteria (1–4). sRNAs are post-transcriptional regulators that bind to multiple target mRNAs, modifying their translation and/or stability (5, 6). In most enterobacteria, the chaperone Hfq plays a critical role in sRNAs regulation by promoting their stability and their interaction with targets mRNA (7, 8). In *Escherichia coli*, over 30 Hfq-dependent sRNAs have been identified, with studies demonstrating their importance in key cellular processes such as biofilm formation, motility, pathogenicity, and stress response (9–11).

Biogenesis of iron-sulfur (Fe-S) cluster gives a striking example of how adaptative responses, coordinated by transcription factors and sRNAs, allow the maintenance of an essential process during adverse conditions. Fe-S clusters, found in both eukaryotes and prokaryotes, play a crucial role as protein cofactors in a wide variety of biological processes such as respiration, metabolism, and genetic regulation (12–15). However, Fe-S clusters are highly sensitive to stress and their biogenesis is therefore tightly regulated. In particular, reactive oxygen species (ROS) such as H_2_O_2_ can destabilize Fe-S clusters through the oxidation of their iron atoms, leading to the inactivation of Fe-S cluster containing proteins (16, 17). In contrast, anaerobic growth is favorable for Fe-S proteins, due to the absence of oxidative stress and the bio-availability of reduced Fe^2+^, which is why this cofactor was probably early adopted during evolution. In fact, recent studies suggest Fe-S clusters biogenesis machineries were present in the last universal common ancestor (LUCA)(12).

In vivo, two multi-protein complexes, Isc and Suf, assemble and deliver Fe-S clusters to client proteins in a wide variety of organisms (18–21). The model bacterium *E. coli* uses both the Isc and Suf systems, encoded respectively by the *iscRSUA-hscAB-fdx-iscX* (hereafter shortened *iscRSUA*) and *sufABCDSE* operons. In this bacterium, Isc is considered as the “housekeeping” machinery and Suf as the stress-responsive system. Indeed, as opposed to Isc, Suf displays the ability to assemble and deliver Fe-S clusters under stress-inducing conditions such as iron starvation or oxidative stress (22, 23). Interestingly, cells operating under the Suf system exhibit lower susceptibility to aminoglycosides (24, 25). This resistance phenotype is due to a reduced capacity of Suf to mature respiratory complexes that generate the proton motive force (pmf) necessary for aminoglycoside entry. It is of primary importance to understand how the transition from one system to another is orchestrated in function of the environmental conditions.

In *E. coli,* expression of Fe-S cluster biogenesis systems is mainly controlled by the transcriptional regulator IscR, itself an [2Fe-2S] cluster containing protein encoded by the first gene of the *iscRSUA* operon (26–28). The holo-form of IscR binds the P*_iscRSUA_* promoter (called type 1 promoter) and represses expression of the whole operon, resulting in a homeostatic modulation of *isc* operon expression according to the cellular demand for Fe-S clusters (26, 28, 29). In contrast, under detrimental conditions for Fe-S cluster biogenesis, IscR accumulates in its apo-form which binds the *suf* promoter (noted type 2 promoter) and activates *suf* expression (30, 31). IscR was shown to be essential to *suf* expression (27). This activation depends on IscR protein accumulation, regardless of its Fe-S maturation state, as both holo and apo-IscR can bind type 2 promoters, including P*_suf_*, implying that IscR synthesis must rapidly increase in stress conditions. IscR also controls other genes not directly related to Fe-S homeostasis and it has notably been shown to be associated to the virulence of numerous bacterial pathogens, thanks to the usage of its Fe-S cluster as a redox sensor (32–35).

In addition to IscR, other global regulators are also involved in the control of *suf* expression in response to stress (36). At the transcriptional level, *suf* expression is also activated by the oxidative stress master regulator, OxyR, during oxidative stress and repressed by Fur, the master regulator of iron homeostasis, when iron is abundant (22, 31, 37). In addition to these transcriptional regulators, the Fur-controlled RyhB sRNA plays a key role in regulating Fe-S cluster biogenesis during iron starvation (38). RyhB base-pairs with the *iscRSUA* mRNA in between *iscR* and *iscS*, leading to RNaseE-dependent degradation of the mRNA 3’-end. The 5’-part coding *iscR* remains stable thanks to the presence of a stabilizing structure that blocks RNaseE action, and IscR is translated. In this way, RyhB was proposed to facilitate the accumulation of apo-IscR during iron starvation and thus to facilitate *suf* expression. Interestingly, RyhB was shown to induce aminoglycosides resistance during iron starvation through its regulation of the Fe-S biogenesis machineries, and to a direct repression of the respiratory complexes (39).

Here, we asked if other sRNAs could regulate Fe-S cluster biogenesis by controlling the expression of the master regulator IscR. To do so, we tested the effect of 26 Hfq-dependent sRNA present in *E. coli* on the *iscR* expression. We found that two of them, OxyS et FnrS, regulate directly *iscR* expression by binding to the 5’-part of the *iscR* mRNA. Interestingly, OxyS and FnrS are expressed in response to opposite conditions and have opposite effects on *iscR* expression. OxyS is synthetized in response to H_2_O_2_ through OxyR activation (40, 41). OxyS was one of the first identified sRNAs in *E. coli* and was described as protecting cells from oxidative stress induced DNA damage by arresting growth under those conditions (42). FnrS is under the control of the FNR regulator and is expressed upon switch to anaerobic growth (43, 44). FnrS represses the expression of multiple genes that are unnecessary under those conditions. Interestingly, *iscR* had already been identified as a potential FnrS target through a bioinformatic approach (45). We further characterize these regulatory processes here and unveil that OxyS promotes resistance to aminoglycosides during oxidative stress through its control of *iscR*.

## Results

### The sRNAs OxyS and FnrS regulate the expression of *iscR* in opposite ways

To investigate the regulation of *iscR* expression by sRNAs, we generated a strain in which the 5’-UTR and the 30 first nucleotides of *iscR* were translationally fused to *lacZ* and cloned downstream of a P_BAD_ promoter, that can be induced by arabinose. This fusion was inserted at the *lac* site on the chromosome of *E. coli* (Fig. 1A). This strain was transformed with a collection of plasmids expressing 26 different *E. coli* Hfq-dependent sRNAs, controlled by a Plac promoter (46). Beta-galactosidase activity was measured after growing cells in a microtiter plate for 6 hours in presence of arabinose and IPTG to induce sRNA expression. Only two sRNAs, OxyS and FnrS, had a significant impact on the P_BAD_-*iscR-lacZ* fusion expression (Fig. 1B). OxyS overexpression induced a 2-fold increase of *iscR* expression, whereas FnrS overexpression resulted in a 16-fold decrease of *iscR* expression, compared to the control strain. To confirm these results, we measured activity of the P_BAD_-*iscR-lacZ* reporter strain transformed either with the control vector plac, pOxyS, or pFnrS, and strains were grown in aerated flasks (Fig. 1C). Overexpression of FnrS had a strong repressing effect on the expression of *iscR,* while OxyS activated its expression by about 2-fold, confirming the results obtained using the screen described above.

**Figure 1:**
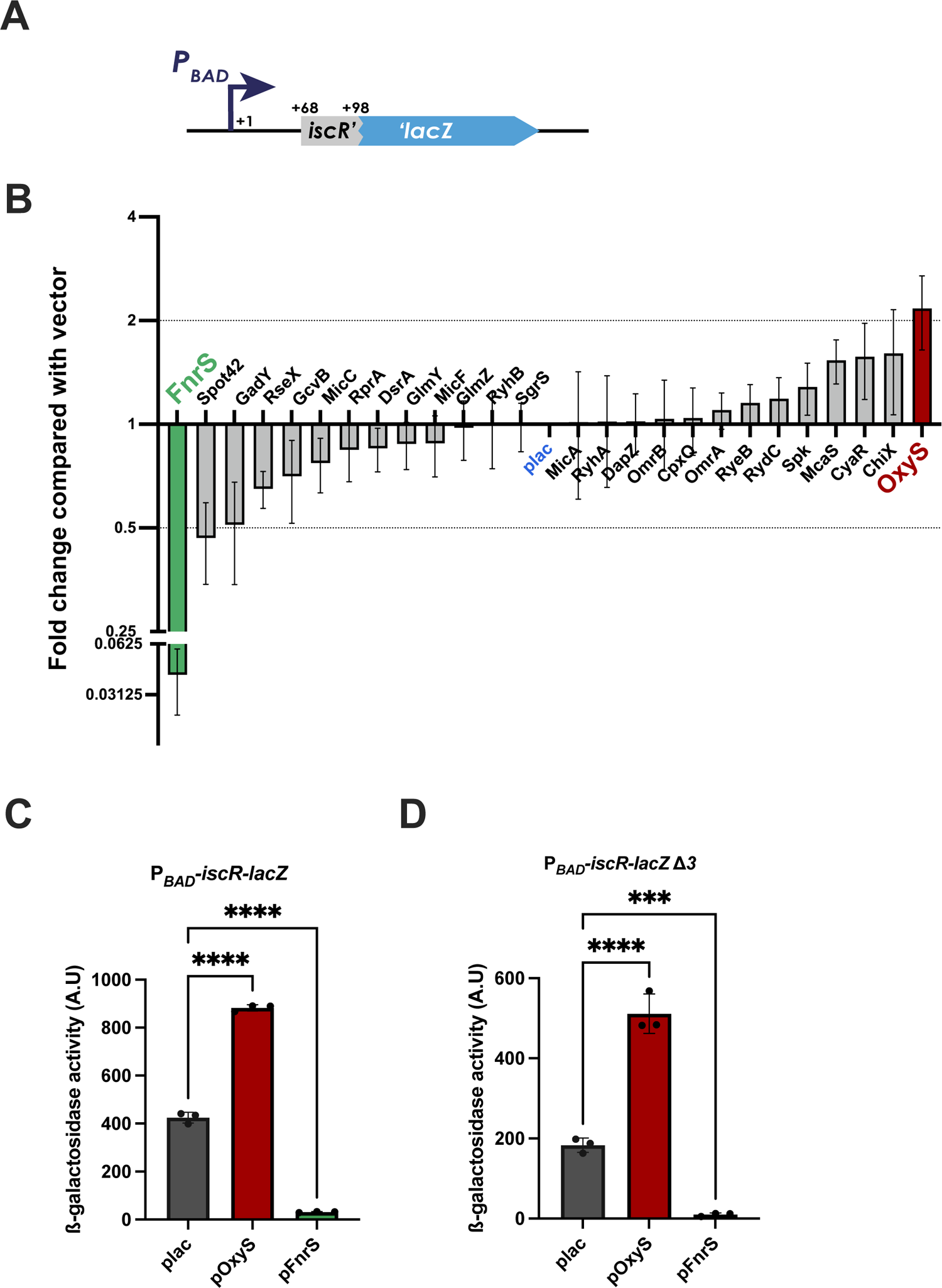
Screening of a sRNA library unveils *iscR* post-transcriptional regulation by two sRNAs. **(A)** Schematic representation of the P_BAD_-*iscR-lacZ* translational fusion inserted at the *lac* site in strain PM3030 used in the sRNA library screening. The transcriptional start site of the *iscR mRNA* (noted +1) was cloned under the control of an arabinose inducible PBAD promoter, and the *iscR* ORF sequence was fused in frame to *lacZ* after the 10^th^ codon (position +98 nt). **(B)** Plasmid allowing the overexpression of 27 sRNAs, as well as the pBR-plac control (plac, in blue) were transformed in strain PM3030 containing the P_BAD_-*iscR-lacZ* fusion. Cells were grown for 6h in presence of 100 µM IPTG before beta-galactosidase activity was measured. Effect of the overexpression of each individual sRNA is represented as the ratio of the effect of each psRNA on the pBR-plac vector. Only fold changes ≥ 2 or ≤ 0.5 were considered significant. Red and green bars indicate sRNAs having an activating or repressing effect, respectively. **(C)** and **(D)** The plasmids pBR-plac (plac, in grey), pOxyS (in red) and pFnrS (green) were transformed in strain PM3030 **(C)** or its isogenic derivative deleted of *oxyS*, *fnrS* and *ryhB* (Δ*3*) (CB433) **(D)**. Cells were grown for 6h in aerated flasks in presence of 100 µM IPTG before beta-galactosidase activity was measured. Error bars represent the standard deviation of at least three independent experiments. Statistical significance was measured by one-way Anova test. ***: P ≤ 0.001; ****: P ≤ 0.0001.

As holo-IscR represses its own expression, we further characterized the action of OxyS and FnrS when *iscR* is under the control of its own promoter. To do so, we constructed a strain containing an *iscR-gfp* fusion under the control P*_iscR_*, integrated chromosomally at the *lac* site (Fig. 2A). Overexpression of OxyS increased expression of the P*_iscR_-iscR-gfp* fusion, while FnrS repressed it, as seen on agar-plates (Fig. 2B), or by following expression over time in cells growing in a microtiter plate, thus demonstrating that OxyS and FnrS conserved their effect even when *iscR* was expressed from its own promoter (Fig. 2C; Fig. S1A and Fig. S2 A and B). It has been previously observed that overexpression of some sRNAs can antagonize the action of others, notably by titrating the Hfq chaperone (47). We thus wanted to confirm that FnrS and OxyS action was independent of each other, as well as of the sRNA *ryhB* that had previously been shown to regulate *iscS* expression (38). To do so, we repeated the previous experiments by overexpressing OxyS and FnrS in the strains containing either the P_BAD_-*iscR-lacZ* fusion (Fig. 1D) or the P*_iscR_-iscR-gfp* fusion, each strain deleted for both *fnrS*, *oxyS* and *ryhB* (Fig. 2D; Fig. S1B and Fig. S2C and D). In each case, effects exerted by OxyS or FnrS were independent of the presence of the other RNA as well as of the presence of *ryhB* (Fig. 2D). Taken together, these data show that FnrS and OxyS act independently of each other and of RyhB, and that they both act post-transcriptionally and in opposite ways on the expression of *iscR*.

**Figure 2:**
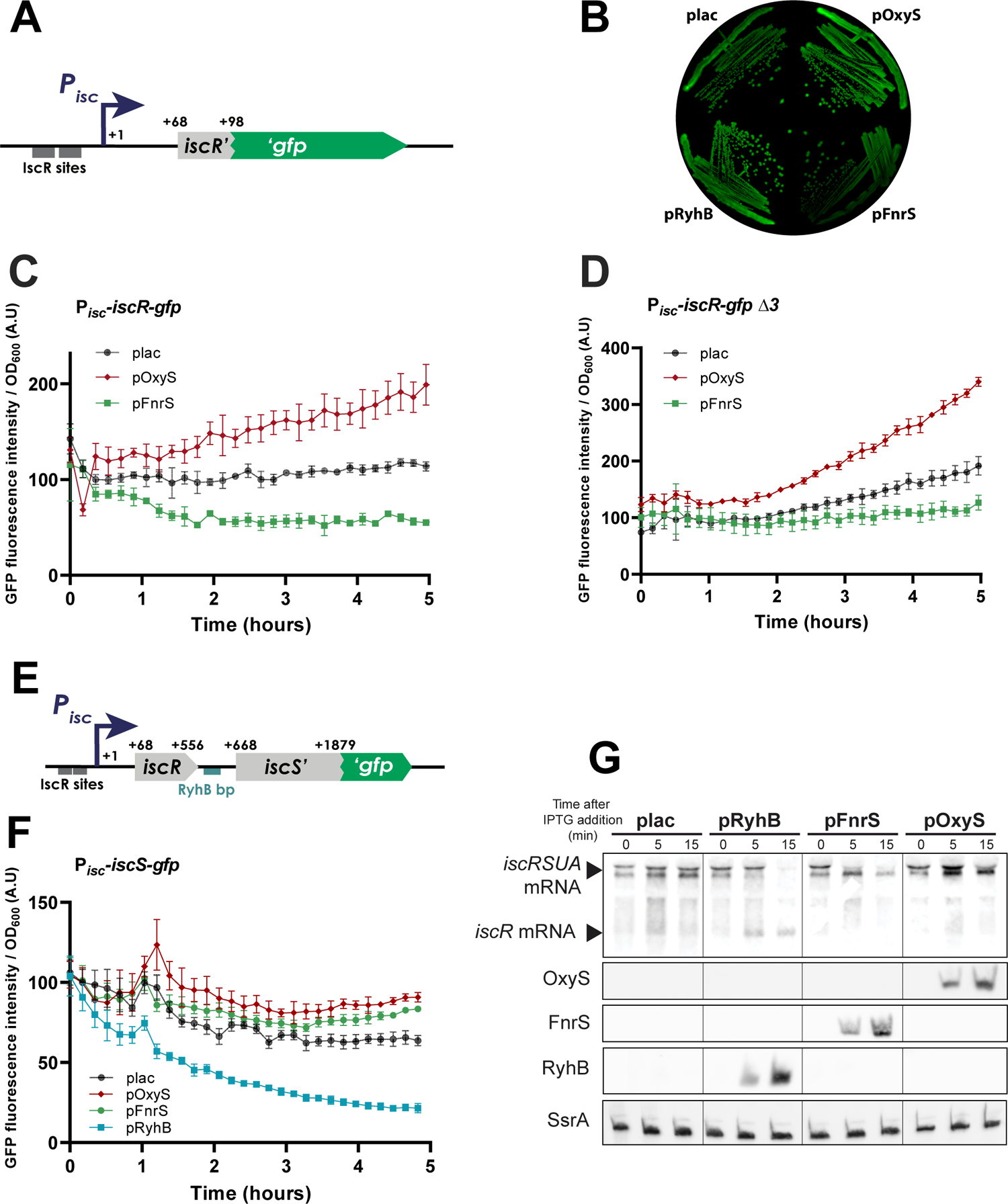
OxyS and FnrS regulate *iscR* expression. **(A)** Schematic representation of the P*_isc_-iscR-gfp* fusion inserted at the *lac* site in strain CB47. This fusion is under the control of the P*_isc_* promoter (dark blue) containing two IscR sites (dark grey). The *iscR* gene from its transcriptional start to its 98^th^ nucleotide (corresponding to its 10^th^ amino acids) is translationally fused in frame to the *gfp* gene. **(B)** The plasmids pBR-plac (plac), pOxyS, pFnrS and pRyhB were transformed in strain CB47 and transformants were streak on a LB agar plate containing 10 µM of IPTG. GFP fluorescence of the colonies was visualized using an Imagequant LAS4000 imager (GE Healthcare). **(C)** and **(D)** OxyS and FnrS overexpression effect on the P_isc_-*iscR-gfp* fusion in culture over time. The plasmids pBR-plac (plac), pOxyS and pFnrS were transformed in strain CB47 (C) or its isogenic derivative deleted of *oxyS*, *fnrS* and *ryhB* (*Δ3*) (CB176) (D). Strains were grown in 96 wells microtiter plates; growth and fluorescence intensity of the strains were measured over time after the induction of sRNAs overexpression by adding 100 µM of IPTG to each culture (growth curves are represented in Fig. S1 A and S1B). A parallel control experiment without adding IPTG was performed in parallel (Fig S2). **(E)** Schematic representation of the P*isc*-*iscS’-gfp* fusion inserted at the *iscS* site in strain CB56. This fusion contains the P*_isc_* promoter (dark blue) and the RyhB basepairing site (light blue) in the *iscR-iscS* intergenic region. Numbers represent positions in nucleotides relative to the *iscRSUA* transcriptional start site. **(F)** The plasmids pBR-plac (plac), pOxyS, pFnrS and pRyhB were transformed in strain CB47 containing the the P*isc*-*iscS’-gfp* fusion. Fluorescence was measured as in (C). Controls showing the growth curves of the different strains and a parrallel experiment without IPTG induction is represented in Fig. S3. **(G)** Northern blot showing the effect of OxyS, FnrS and RyhB overexpression on the *iscRSUA* and *iscR* mRNA. The plasmids pBR-plac (plac), pOxyS, pFnrS and pRyhB were transformed in strain PM1490. Strains were grown in LB to OD_600_ = 0.6 at which point 100 µM IPTG was added to the cultures. Sampling has been performed immediately before (0), 5 and 15 minutes after the induction of sRNAs overexpression using 100 µM IPTG in the PM1490 strain. Error bars represent the standard deviation of at least three independent experiments.

### OxyS and FnrS have no effect on the rest of the *isc* operon

To further characterize the regulatory role of the sRNAs on the rest of the *iscRSUA* operon, we constructed an *iscS-gfp* fusion at the endogenous *isc* site (Fig. 2E). Of note, this strain did not exhibit any visible growth defect (as opposed to known growth defects of *iscS* deletion), indicating that the resulting protein fusion is functional (Fig. S3 A-C). We then tested expression of the *iscS-gfp* fusion over time after overexpressing OxyS and FnrS from their respective plasmids (Fig. 2F). As a control, we also performed the same experiment in a strain overexpressing the RyhB sRNA which was previously shown to induce degradation of the 3’-part of the *iscRSUA* mRNA, inducing the accumulation of a 5’ mRNA fragment containing *iscR* (38). While RyhB effectively drastically inhibited the expression of *iscS* when overexpressed, expression of OxyS or FnrS had very modest effects on the expression of *iscS-gfp* as compared to the effect on *iscR* (Fig. 2F). These results indicate that OxyS and FnrS mainly act on the expression of *iscR* without affecting that of the rest of the operon. In addition, while overexpression of RyhB induced the degradation of the full length *iscRSUA* mRNA, it was still detected when either OxyS or FnrS were overexpressed (Fig. 2G). We note that, in agreement with their effect on *iscR* expression, OxyS induced a slight increase of the *iscRSUA* mRNA levels while FnrS induced a slight diminution of the same mRNA. Taken together, the fact that OxyS and FnrS do not influence drastically *iscS* expression and *iscRSUA* mRNA levels suggest that they exert their control mainly translationally, most likely by preventing or promoting ribosome access to the *iscR* mRNA RBS.

### OxyS and FnrS regulate *iscR* expression by binding to the *iscR mRNA* 5’UTR

We used the *in-silico* structure prediction program Mfold to investigate the potential base-pairing and mode of action of the sRNAs on the *iscR mRNA* (48). This analysis showed that the 5’ part of the *iscR* mRNA could form a stem-loop structure which may partially block the access of the ribosome to its binding site (Fig. 3A). FnrS has previously been predicted and then experimentally confirmed to base-pair at the level the ribosome binding site (RBS) of the *iscR* mRNA, thus likely inhibiting ribosome access at this site, which is consistent with its repressing activity on *iscR* expression we observed (Fig. 3B) (45). Interestingly, using Mfold, the OxyS sRNA was predicted to base-pair on the 5’-part of this stem loop structure, which could induce its opening and release the access of the RBS for the ribosome, consistent with an activating role of OxyS in the translation of *iscR* (Fig. 3C).

**Figure 3:**
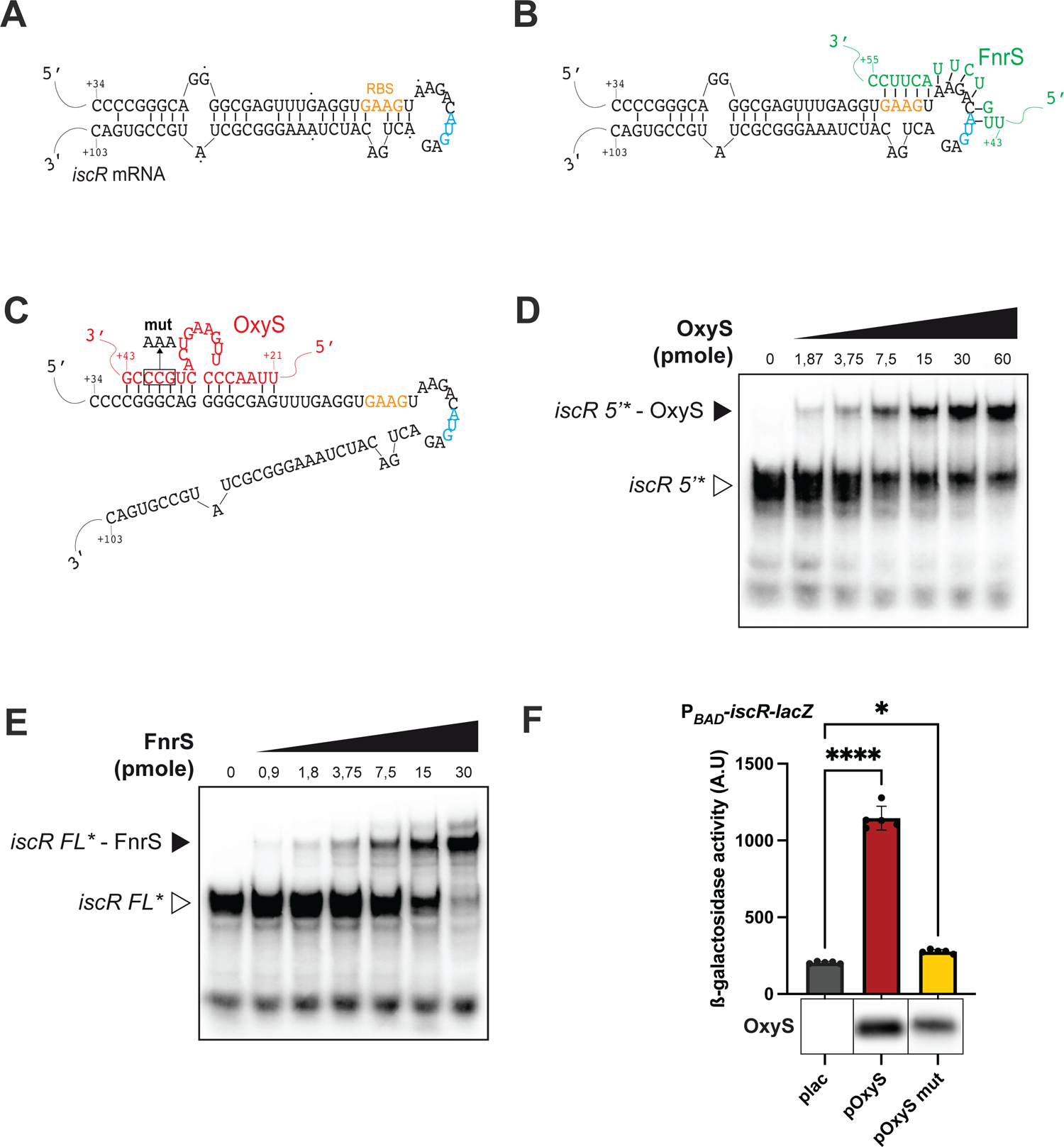
OxyS and FnrS post-transcriptionally regulate *iscR* by base-pairing on its stem-loop. **(A)** Secondary structure prediction of the *iscR* mRNA 5’ UTR using the Mfold program (48). The putative RBS is represented in orange, the AUG translation start is colored in blue. **(B)** Base-pairing prediction of FnrS (green) on the *iscR* mRNA using Mfold. Of note, this base-pairing was also predicted by (45) **(C)**. Base-pairing prediction of OxyS (red) base-pairing using Mfold. In each case, numbers represent positions on the mRNA relative to the transcriptional start site of *iscR* or of the sRNAs. **(D)** Native EMSA of ^32^P-labeled *iscR* 5’ (0.3 pmole) and increasing amounts of OxyS. **(E)** Native EMSA of ^32^P-labeled *iscR* full length (FL) (0.3 pmole) with increasing amounts of FnrS. **(F)** Overexpression of a mutated allele of OxyS, OxyS mut (GCC -> AAA) as represented in (C), on the P_BAD_-*iscR-lacZ* translational fusion (PM3030) compared to pOxyS and empty vector. The levels of the OxyS WT or mutated sRNA were measured above the graph using Northern blot from samples of one representative experiment. Error bars represent the standard deviation of three independent experiments. Statistical significance was measured by one-way Anova test. *: P ≤ 0.05; ****: P ≤ 0.0001.

We performed electrophoretic mobility shift assays (EMSAs) to test the base-pairing of OxyS with the *iscR* mRNA. A clear band shift was observed when the sRNA was incubated with a radio-labelled 5’-UTR of the *iscR* mRNA (from nucleotides 1 to 82) comprising only the 5’-part of the stem (Fig. 3D). A shifting duplex was also observed when OxyS was incubated with a larger version of the 5’-part of the *iscR* mRNA comprising the full stem loop predicted structure, albeit in a less efficient way, indicating that binding of the sRNA may be less efficient when the stem loop is formed and/or in absence of Hfq (Fig. S4A). FnrS was also able to efficiently form a duplex with the radio labelled 5’-part of the mRNA (Fig. 3E).

To further confirm the base-paring of OxyS to the *iscR* mRNA in vivo, we mutated three nucleotides in the OxyS sequence involved in the predicted base-paring (OxyS mut) (Fig. 3C). Overexpression of this mutated form of OxyS was not able to activate the expression of a P_BAD_-*iscR-lacZ* fusion as compared to the OxyS WT (Fig. 3F). Furthermore, in vitro, the mutated version of OxyS was not able to induce a shift of the *iscR* mRNA as seen by EMSA (Fig. S4B). Taken together, these results show that both sRNAs bind directly the *iscR* mRNA to exert their regulation, most likely by inhibiting ribosome access for FnrS and oppositely for OxyS.

### FnrS regulation of *iscR* during anaerobiosis

To validate the role of FnrS in physiological conditions, we tested the ability of the sRNA to regulate *iscR* during anaerobic growth. To do so, the strain containing an P_BAD_-*iscR-lacZ* fusion (Fig. 1A), was deleted or not for *fnrS* and grown anaerobically in Hungate’s tube before beta-galactosidase assays were performed. We first verified by Northern blots that FnrS was indeed expressed in these conditions (Fig. S5-A). The expression of the P_BAD_*-iscR-lacZ* fusion was drastically enhanced in absence of *fnrS*, further validating the role of the sRNA in vivo conditions (Figure S5-B). As anaerobic conditions are favorable for Fe-S biogenesis, we hypothesize that FnrS, by inhibiting *iscR* production, spares Fe-S biogenesis and blocks unnecessary production of Suf in these conditions.

### OxyS activates *iscR* expression during oxidative stress

The expression of *oxyS* is up-regulated by OxyR in presence of H_2_O_2_ oxidative stress (40). In these conditions, and in slight contrast to anaerobiosis, ROS destabilize Fe-S clusters and induce a change in Fe-S biogenesis machinery usage (22). OxyR also strongly induces an arsenal of catalases (*katE* and *katG*) and peroxidase (*ahpCF*) to quickly detoxify the medium of H_2_O_2_ (49). In order to induce a protracted oxidative stress to assess the effect of OxyS on *iscR* expression in stress conditions, we used a strain named Hpx^-^ that is devoid of catalases and peroxidases. This strain can be grown normally and manipulated genetically in anaerobic conditions. When switched to aerobic growth, the Hpx^-^ strain shows an accumulation of intracellular H_2_O_2_ (up to 1 µM), which has previously been shown to activate the expression of the OxyR regulon (49). As expected, we see a protracted accumulation of OxyS over time when the Hpx^-^ strain is switched from anaerobic to aerobic growth, which is not the case for the control Hpx^-^ Δ*oxyS* strain (Fig. 4A).

**Figure 4:**
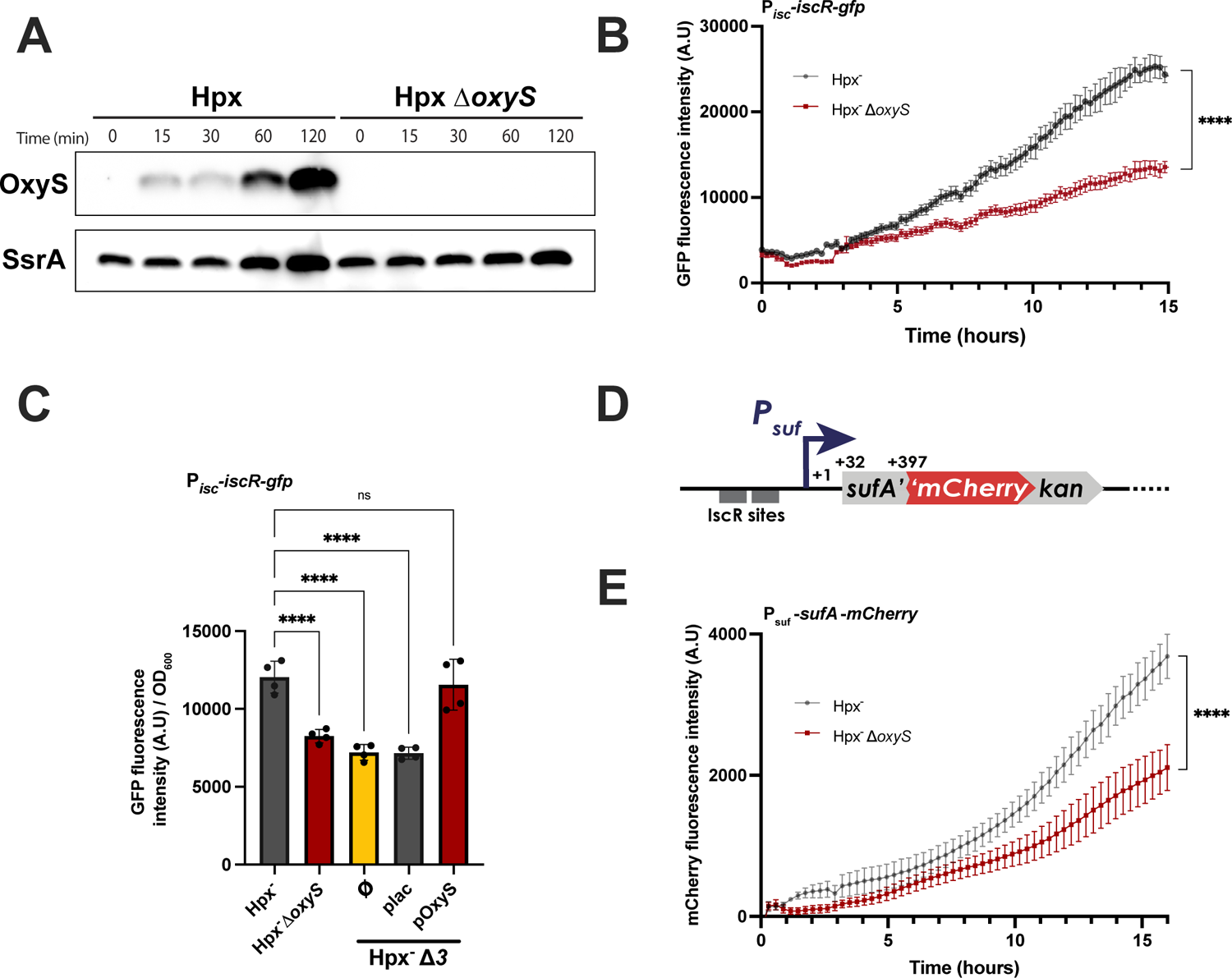
Effect of OxyS on *isc* and *suf* expression under oxidative stress. **(A)** Northern blot showing OxyS expression after the induction of H_2_O_2_ oxidative stress. The Hpx^-^ strain and its isogenic derivative deleted for *oxyS* (CB178) were grown anaerobically up to OD_600_ :: 0.3 (0 min) and then transferred aerobically leading to oxidative stress induction. Samples were taken over time and Northern blots were performed using probes agains OxyS or the SsrA RNA as a control. **(B)** *iscR* expression followed by fluorescence in the presence (grey) or absence of *oxyS* (red) after oxidative stress induction. The Hpx-strain containing the P_isc_-*iscR-gfp* fusion (CB182) and its isogenic derivative deleted for *oxyS* (CB186) were cultured anaerobically up to OD_600_ :: 0.3 and then transferred aerobically to induce oxidative stress. Fluorescence intensity was measured every 15 minutes during 15 hours after oxidative stress induction. Growth of the strains was measured in parallel and was identical (Fig. S6). **(C)** Expression of the P_isc_-*iscR-gfp* fusion inserted in the Hpx^-^ strain (CB182) or its isogenic derivative deleted for *oxyS* (CB186) or for *oxyS*, *ryhB* and *fnrS* (CB203). The strain CB203 was further transformed with plac or pOxyS to complement the *iscR* fusion. The fluorescence intensity was measured 4 hours after oxidative stress / *oxyS* expression induction. **(D)** Schematic representation of the *sufA-mCherry* fusion inserted at the endogenous *suf* site in the Hpx-strain (strain CB190). This fusion is under the control of the P*_suf_* promoter (dark blue) containing two IscR sites (dark grey). The *sufA* gene is translationally fused in frame to the *gfp* gene. (**E)** *sufA* expression during oxidative stress in Hpx^-^ containing the *sufA-mCherry* fusion (CB190) and its isogenic Δ*oxyS* derivative (CB200). Strains were grown anaerobically up to OD_600_ :: 0.3 (0 min) and then transferred aerobically leading to oxidative stress induction. Fluorescence intensity was measured every 15 minutes during 16 hours after oxidative stress induction. Growth of the strains was measured in parallel and was identical (Fig. S8). Error bars represent the standard deviation of at least three independent experiments. Statistical significance was measured by one-way Anova test. ****: P ≤ 0.0001.

To test the role of OxyS on *iscR* expression during oxidative stress, we introduced the P_isc_-*iscR-gfp* translational fusion (Fig. 2A) in the Hpx^-^ strain deleted or not for *oxyS* and followed expression of the fusion over time after switching the cells from anaerobic to aerobic, oxidative stress inducing, conditions (Fig. 4B; S6). Strikingly, P_isc_-*iscR-gfp* expression was turned on as soon as the cells were grown in presence of oxygen in the Hpx^-^ *oxyS^+^* strain and reached its maximum of expression after several hours of exposition to oxidative stress. In contrast, *iscR* expression was twice as less important in the isogenic strain devoid of *oxyS* (Fig. 4B and Fig. S7 A-B). A similar effect was seen in an Hpx^-^ strain deleted of *oxyS*, *fnrS* and *ryhB* (Hpx^-^ Δ*3*), indicating that the other sRNAs known for their role in Fe-S cluster biogenesis regulation did not play a role on *iscR* expression in these conditions (Fig. 4C). In agreement with the direct role of OxyS, plasmidic overexpression of OxyS in the Hpx^-^ Δ*3* complemented this phenotype by restoring expression of the P_isc_-*iscR-gfp* fusion to the same levels as the Hpx^-^ (Fig. 4C). Taken together, these results demonstrate that OxyS is necessary for *iscR* optimal expression in response to oxidative stress.

### IscR activation by OxyS enhances *suf* expression during oxidative stress

Sub-micromolar concentrations of H_2_O_2,_ such as the ones encountered in the Hpx^-^ strain during aerobic growth, have been shown to poison the Isc machinery and to induce *suf* expression to compensate for a deficiency in Fe-S cluster biogenesis (16, 22). Oxidative stress induction of *suf* expression has been previously shown to be mediated by OxyR and accumulation of apo-IscR (27, 37). We thus asked if OxyS could play a role in the induction of Suf through its activation of *iscR* expression. To test this hypothesis, we constructed Hpx^-^ strains containing a P*_suf_*-*sufA-mCherry* fusion at its endogenous locus, with or without *oxyS*, and measured fluorescence intensity of the strain after switching the cultures from anaerobic to aerobic conditions (Fig. 4D-E; Fig. S8). As expected, oxidative stress induction turned on *suf* expression in the OxyS containing cells and reached a peak after several hours of growth. In the *oxyS* deficient strain, induction was slower and remained lower (about 2-fold) during the course of the growth, thus showing that OxyS is necessary for a full expression of *suf* in oxidative stress conditions.

### OxyS induces resistance to aminoglycosides during oxidative stress through its control of Fe-S biogenesis

Usage of the Suf machinery in stressful conditions has been previously associated with a transient resistance to aminoglycosides (24). In fact, we have previously shown that RyhB regulation of the Isc / Suf balance during iron starvation is responsible for a resistance phenotype to gentamicin, a major antibiotic of the aminoglycosides class, when iron is limiting (39). We thus tested whether OxyS could be responsible for a similar phenomenon during oxidative stress.

We first assayed if oxidative stress could modulate *E. coli* sensitivity to various classes of antibiotics by measuring the minimum inhibitory concentrations (MIC) of antibiotics in the Hpx^-^ strain grown in aerobic conditions, and thus accumulating sub-micromolar concentrations of H_2_O_2_. Sensitivity of the Hpx^-^ strain to ampicillin was unaffected as compared to the WT strain; the Hpx^-^ strain was slightly more sensitive than the WT to norfloxacin and tetracycline. In each of those cases, deleting *oxyS* from the Hpx^-^ strain did not affect sensitivity to these antibiotics, indicating that the sRNA is not involved in these phenotypes (Fig. 5A).

**Figure 5:**
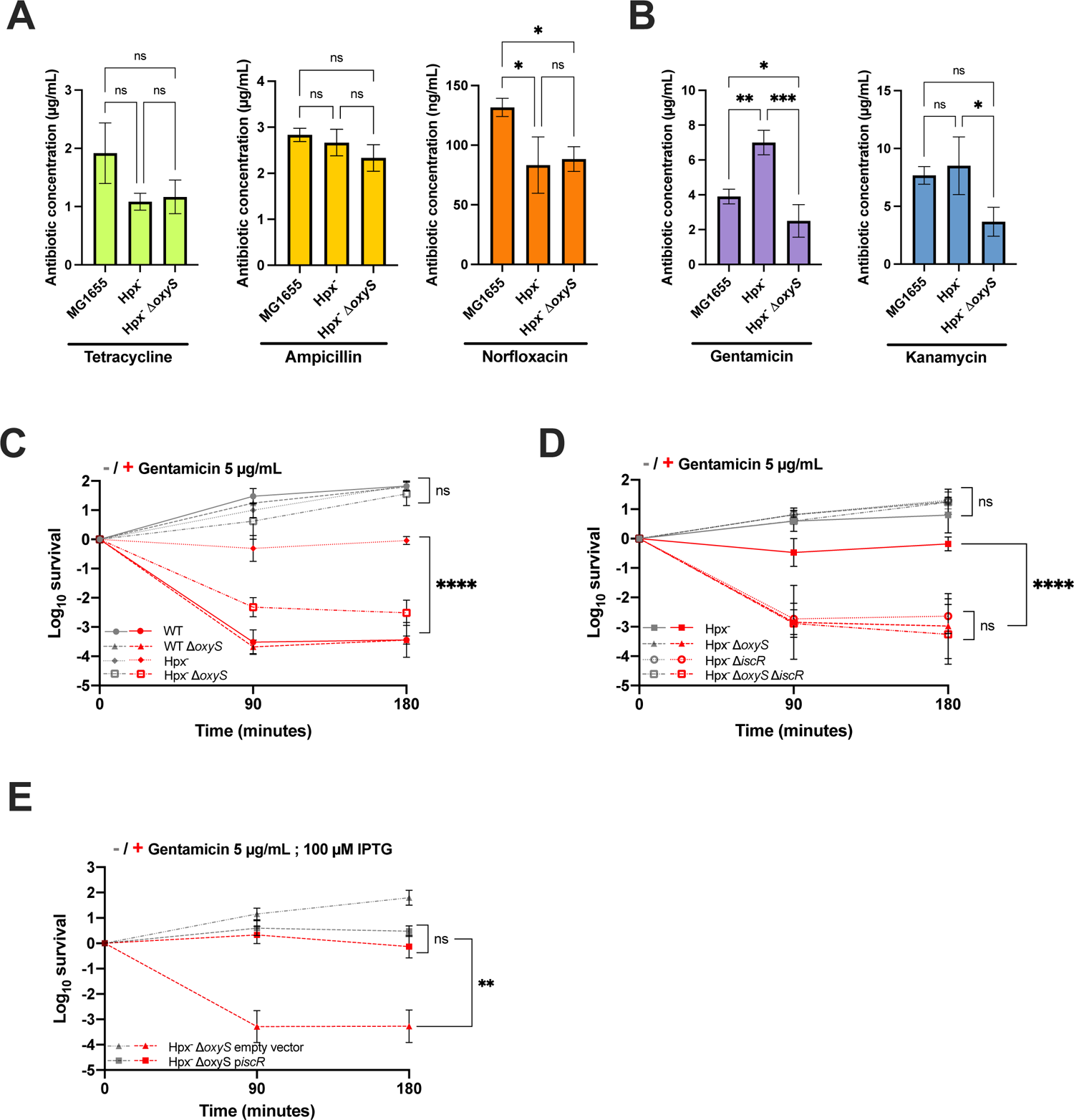
H_2_O_2_ oxidative stress induces aminoglycosides resistance in *E. coli* trough OxyS regulation of *iscR* expression. **(A)** Minimal Inhibitory Concentration (MIC) of several antibiotic classes. MIC of tetracycline (green), ampicillin (yellow) and norfloxacin (orange) were determined for MG1655, Hpx^-^ and Hpx^-^ Δ*oxyS* (CB178) strains under aerobic conditions **(B)** MIC values of aminoglycosides (in µg/mL). MIC of Gentamicin (purple) and kanamycin (blue) were determined for the same strains as in (A). In (A) and (B) MIC are represented as an average of three individual experiments. **(C)** Survival of MG1655, MG1655 Δ*oxyS* (CB8), Hpx^-^ and Hpx^-^ Δ*oxyS* (CB178) after gentamicin (5 µg/mL) treatment (in red). Untreated cultures are represented in grey. During the experiment, only the Hpx^-^ strains are experiencing H_2_O_2_ oxidative stress. **(D)** Survival of Hpx^-^, Hpx^-^ Δ*oxyS* (CB178), Hpx^-^ Δ*iscR* (CB226) and Hpx^-^ Δ*oxyS* Δ*iscR* (CB228) were determined with or without gentamicin (5 µg/mL) treatment (in red). **(E)** Survival of Hpx^-^ Δ*oxyS* strains transformed with p*iscR* (squares) or with the empty vector (triangles). The strains were cultivated with 100 µM of IPTG to induce *iscR* overexpression. Colony forming units (CFU) were determined by counting the number of surviving bacteria after 90 or 180 minutes of growth in presence or not of gentamicin. Each value was normalized to T0 and plotted as log_10_ of surviving bacteria. CFU at T0 was approximatively 1.5 x 10^7^ CFU / mL for each sample. Error bars represent the standard deviation of three independent experiments. Statistical significance was measured by one-way Anova test. ** P ≤ 0.01; ****: P ≤ 0.0001.

Interestingly, the Hpx^-^ strain was more resistant to the aminoglycoside gentamicin than the WT strain (doubling of the MIC, Fig. 5B). This resistance phenotype was lost when *oxyS* was deleted from this strain. Furthermore, while the MIC of the Hpx^-^ strain was comparable to that of the WT strain for another aminoglycoside, kanamycin, the Hpx^-^ Δ*oxyS* strain was twice as more sensitive to this antibiotic (Fig. 5B). To further characterize this phenotype, we measured the survival of the WT and Hpx^-^ strains grown aerobically in presence of gentamicin (5 µg/mL). WT cells, that do not experience oxidative stress, were rapidly cleared from the medium at these antibiotic concentrations, regardless of the presence of OxyS (Fig. 5C). In contrast, the Hpx^-^ strain exhibited a strong resistance to gentamicin at this concentration (no killing). Interestingly, this resistance was completely lost when *oxyS* was deleted from the Hpx^-^ strain. Taken together, these results indicated that oxidative stress antagonized aminoglycosides in an OxyS dependent manner.

We reasoned that if OxyS renders Hpx^-^ cells resistant to aminoglycosides through its positive control of *iscR*, an Hpx^-^ strain devoid of *iscR* should be sensitive to aminoglycosides. This is indeed the phenotype we observed when Hpx^-^ Δ*iscR* cells were grown in presence of gentamicin (Fig. 5D). In addition, further deleting *oxyS* in the Hpx^-^ Δ*iscR* did not sensitize this strain to gentamicin, suggesting that other targets of OxyS do not participate in the resistance phenotype of the Hpx^-^. Reciprocally, over-expressing *iscR* devoid of the OxyS base-pairing site from a plasmid was sufficient to fully restore the gentamicin resistance of an Hpx^-^ Δ*oxyS* strain (Fig. 5E). Taken together these results demonstrate that OxyS induces resistance to aminoglycosides during oxidative stress, through its positive control of IscR.

## Discussion

Using a library of plasmids enabling the overexpression of 26 Hfq-dependent sRNAs, we have here uncovered that two sRNAs, FnrS and OxyS, control *iscR* expression. Remarkably, these sRNAs act in opposite ways and in response to opposite conditions: OxyS, expressed during oxidative stress, increases the expression of *iscR*, while FnrS, expressed during anaerobiosis, represses it. With the addition of RyhB, which has previously been found to repress the expression of the Isc machinery and of the *erpA* Fe-S cluster transporter during iron starvation (38, 50), our study thus shows that at least three sRNAs regulate Fe-S clusters biogenesis, highlighting the importance of finely regulating this process in the cell. Interestingly, all three sRNAs act in function of environmental conditions that are deeply connected to Fe-S clusters homeostasis: iron starvation and oxidative stress are well known perturbators of Fe-S clusters homeostasis and biogenesis, while anaerobiosis can be considered as the most favorable condition for biogenesis of Fe-S clusters due to absence of ROS and availability of reduced iron (Fe^2+^).

Based on our results, we propose a model for how OxyS and FnrS control Fe-S clusters biogenesis through their regulation of IscR (Fig. 6). In the case of OxyS, upon oxidative stress induction, OxyR oxidation will result in OxyS production, which will in turn activate IscR translation post-transcriptionally. In those conditions, newly synthetized IscR will accumulate in its apo-form and bind to type 2 promoters such as the *suf* promoter (22). Thus, by activating expression of *iscR*, OxyS indirectly activates *suf* expression to maintain Fe-S cluster homeostasis. As an indirect consequence, this switch in Fe-S biogenesis systems reduces sensitivity to aminoglycosides, most likely because respiratory complexes are inefficiently matured by the Suf system, thus reducing aminoglycosides entry.

**Figure 6:**
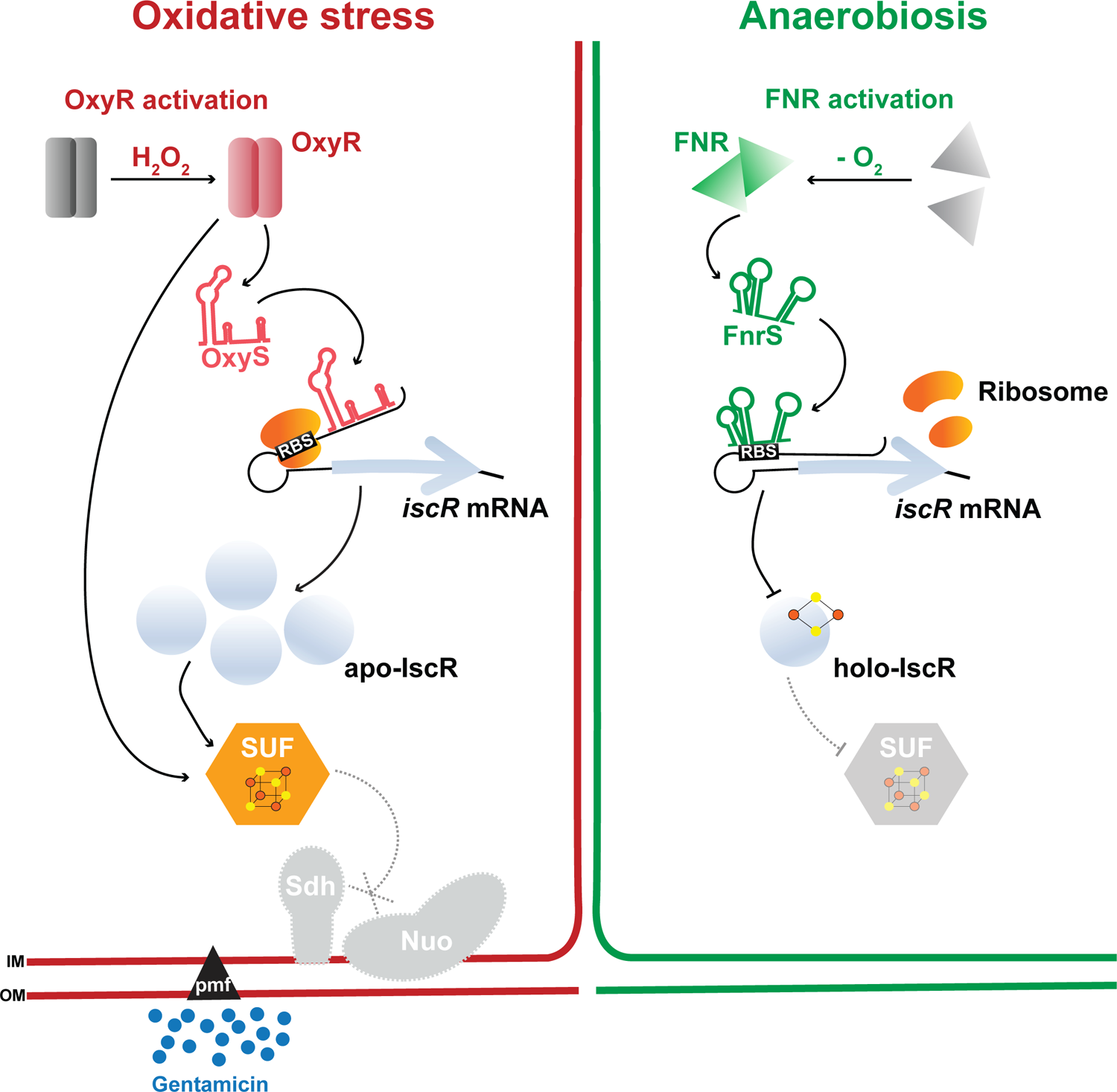
Model for the regulation of Fe-S biogenesis by OxyS and FnrS. The presence of H_2_O_2_ leads to OxyR activation (left panel) and thus of the expression of the OxyR regulon to which the sRNA OxyS belongs. Through its base-pairing to the potential stem-loop structure formed in the 5’-UTR of the *iscR* mRNA, OxyS increases IscR translation. IscR accumulates in its apo-IscR form and activates *suf* expression in conjunction with the direct activation by OxyR. As the Suf machinery is not able to fully maturates Nuo and Sdh respiratory complexes, the pmf is lowered, leading to a diminished gentamicin entry inducing resistance. Conversely, in absence of oxygen (right panel), FNR activates the expression of the sRNA FnrS, which binds on the *iscR* RBS and thus inhibits its translation. During anaerobiosis, FnrS by repressing *iscR* expression locks the expression of *suf*, dispensable in those conditions and promotes the use of the Isc machinery.

OxyR, the master regulator of the H_2_O_2_ response that also controls *oxyS* expression, has also been shown to induce *suf* expression upon oxidative stress by directly binding to the *suf* promoter (37). Thus, OxyR, OxyS and IscR participate in a feed forward loop to fire the expression of *suf* after exposure to H_2_O_2_ (22). Mixed “feed-forward loop” involving small RNAs and transcriptional regulators acting on the expression of the same genes / pathways is a common trend for sRNA regulation (51–53). The current hypothesis is that sRNAs provide a double layered regulation to those regulatory motifs, serving to fine tune protein expression and/or act as a noise filtering mechanism. In the present case, while *suf* is expressed upon oxidative stress exposure even in the absence of OxyS, the sRNA is needed for the full activation of the system, reinforcing the importance of Suf to provide Fe-S clusters in such conditions. Absence of this second layer of regulation can have dramatic phenotypic consequences for the bacteria, as evidenced here by the fact that OxyS indirect regulation of Suf drastically decreases sensitivity of bacteria to aminoglycosides.

The OxyS induced aminoglycosides resistance mechanism described here might play a role in the context of infection. Indeed, phagocytes produce superoxide that spontaneously dismutates to H_2_O_2_ in the range of 1 - 4 µM to suppress bacterial growth (54, 55). These doses will activate the OxyR regulon and thus should paradoxically protect bacteria from gentamicin (56). Perhaps even more relevant, H_2_O_2_ has been proposed to accumulate in the lumen of the intestine where *E. coli* and other enterobacteria reside, notably via the production of hydrogen sulfide by sulfate reducing bacteria that will react with oxygen diffusing through epithelial cells (54).

Surprisingly, only few direct targets of OxyS were identified up to now, including the transcription factors *fhlA* and *flhDC*, and more recently the termination factor *nusG* (40, 42, 57). In this last case, it was shown that OxyS overexpression is toxic to the cells because the sRNA indirectly induces cell cycle arrest to allow DNA repair. We propose that the difficulty in uncovering the physiological role of OxyS might be due to the fact that controlling expression of this sRNA beyond toxic levels is difficult with overexpression plasmids or with high doses of external H_2_O_2_. As we show here, this problem may be bypassed using the Hpx^-^ strain which allows for a protracted expression of *oxyS* without apparent toxic effects. Further examining the role of OxyS regulation in such contexts will shed new light on the function of this sRNA in vivo. In a striking contrast to OxyS, FnrS is induced upon anaerobic growth conditions and will precisely have the opposite effect on *iscR* expression by inhibiting its translation. It has before been shown that IscR is essential to expression of *suf* (27). We thus propose that FnrS repression acts as a lock that impedes *suf* expression during anaerobiosis. Furthermore, FnrS overexpression had no visible effect on the expression of the rest of the *isc* operon, encoding the Isc machinery. We thus propose that by inhibiting IscR, FnrS strongly favorizes Isc usage over Suf, both by removing IscR negative retro-control on *isc* and by preventing accumulation of IscR that would drive *suf* expression. In this way, FnrS promotes the expression of the Isc housekeeping system in conditions known to be favorable for Isc function.

Our findings demonstrate that both sRNAs engage in base-pairing with the *iscR* mRNA 5’-UTR, at sites in close proximity to each other (Fig. 4). While a computational study previously identified the base-pairing of FnrS with the *iscR* mRNA (45), the binding of OxyS is a novel finding. Interestingly, both sRNAs were found paired to the *iscR* mRNA in the Ril-seq study conducted by Melamed *et al*. in which the global profile of sRNA-mRNA pairs bound to Hfq were identified but had not been validated in this particular case (58). We could not detect drastic modifications to *iscR* mRNA levels as seen by northern blots, strongly suggesting that the sRNAs act primarily on the translation level. In agreement with this, overexpressing either FnrS or OxyS had modest effects on the expression of the downstream *iscS* gene. This uncoupling between *iscR* and the rest of the operon by FnrS and OxyS is reminiscent of the one that is provided by RyhB regulation of *iscRSUA*. In this last case, RyhB base-pairing leads to the degradation of the 3’ part of the mRNA, encoding the Isc machinery, while the 5’ part containing *iscR* is stabilized and available for translation (38).

The genes encoding IscR, Isc, Suf and the sRNAs OxyS and FnrS are conserved in many enterobacteria and notably in pathogenic species such as *Salmonella*, *Klebsiella*, *Yersinia* or *Pseudomonas* (Figure S9). Strikingly, IscR has been recently shown to be a regulator of the virulence of these bacteria as well as in Enterotoxigenic *E. coli* (34, 59, 60). In these bacteria, IscR is used as an iron availability and/or oxygen sensor to control the expression of virulence genes (32). For instance, in *S. enterica,* IscR has been shown to repress the expression of *hilD*, which encodes the master regulator of the Spi1 pathogenicity highland. Consequently, a *S. enterica* Δ*iscR* mutant is hyper invasive in epithelial cells (34). In *P. aeruginosa,* IscR does not directly regulate the expression of virulence factors, but it is implicated in controlling oxidative stress resistance genes and fimbriae production (61, 62). It is thus likely that regulation of *iscR* expression by OxyS and FnrS well be conserved in other pathogenic bacteria, allowing for a fine tuning of the expression of the virulence genes in function of the oxygen tension and reactive oxygen species found during the infectious process. Further work will be required to fully unravel the potential role of these sRNAs in the virulence process of these bacteria through their control of IscR.

## Materials and Methods

### Strains and cultures

All strains used in this study are derivatives of *E. coli* MG1655 and are listed in Table 1. Strains were grown in LB broth (Difco). Anaerobic cultures were performed in gas tight Hungate tubes containing LB medium deoxygenated by N_2_ bubbling for 20 min before autoclaving. Hungate tubes were inoculated through the septum (1/100 dilution) and strains were grown at 37 °C without shaking. For oxidative stress induction, starting from a 1/100 dilution of an anaerobic overnight culture, the Hpx^-^ strain and its isogenic derivatives were grown anaerobically in Hungate tubes to OD_600_ :: 0.3. The cultures were then transferred in flasks and cultivated aerobically at 37 °C with shaking.

### Genetic constructions

Transductions with P1 phage were used for moving marked fusions and mutations as described previously (63). The different plasmids used in this study are described in Table 1 and have been transformed as previously described in (64). All oligonucleotides used are listed in Table 2. The strain containing the P_BAD_-*iscR-*lacZ fusion (PM3030) and the P_isc_-*iscR-lacZ* fusion (PM2250) were constructed as described in (64) by recombineering in the strain PM1205 with the PCR product obtained by amplifying *iscR* from the MG1655 strain with primer pair P_BAD_-iscR-F and lacZ-iscR-R or Pisc-F and lacZ-iscR-R, respectively. The strain containing the P_isc_-*iscR-gfp* fusion (CB47) was constructed as follows. First, the *gfp-ka*n sequence was amplified by PCR using the pGBMKn-GFP-plasmid as a template and oligonucleotides iscR-GFP-F and lacZ-GFP-Kan-R (see Table 2). Then the purified PCR product were electroporated in the PM2250 strain containing pkD46 plasmid allowing homologous recombination as described in (65). The strain was then curated of the *kan* resistance gene using pCP20 plasmid as previously described in (65). The strains containing the *iscS’-gfp* and *sufA-mCherry* fusions were constructed using recombineering as follows: first, the *gfp-kan* and *mCherry-kan* sequences were amplified with appropriates oligonucleotides couple iscS-GFP-F / iscS-GFP-R and sufA-mcherry-F / sufA-mcherry-R, respectively, and using pGBMKn-GFP and pGBMKn-mCherry plasmids as templates; the PCR products were then introduced on the chromosome of the PM1490 strain using recombineering as previously described. The plasmid pTrc-*iscR* was constructed as follows. First, the coding region of *iscR* from the *E. coli* MG1655 chromosomal DNA was amplified by PCR using the following primers pair: NcoI-iscRup / iscRdown-BamHI (Table 2). The PCR product was then digested by *Nco*I and *Bam*HI and cloned into the *Nco*I and *Bam*HI linearized pTrc99a vector (66). The insert fragment was then verified by DNA sequencing.

### Library screen

The library screen was performed as previously described (64). Briefly, CaCl_2_ transformation technique was performed in microtiter plates, following by spotting cells for selection of transformants on LB ampicillin plates. These spots were then used to inoculates media for overnight growth in microtiter plate. Cultures for ß-galactosidase activity assay were performed by 1/100 dilution of those overnight cultures in microtiter plates. Cultures were grown for approximatively 7 hours at 37°C with shaking before being lysed and ß-galactosidase activity determined as below.

### ß-galactosidase assay

Beta galactosidase assays were performed as in (64). Overnight cultures of strains were diluted 1/100 times in fresh LB medium containing ampicillin and IPTG (isopropyl ß-D-1-thiogalactopyranoside). After approximatively 7 hours of growth, 100 µL of the cultures were transferred to microtiter plates, in triplicates for each condition. The absorbance at 600 nm was measured using a microtiter plate reader (Spark 10M, Tecan). Next, 50 µL of permeabilization buffer (100 mM Tris HCl pH 7.8; 32 mM Na_2_HPO_4_; 8 mM EDTA; 40 mM Triton) and the microtiter plate was incubated during 10 minutes at RT. Then, O-Nitrophenyl-ß-D-galactopyranoside (ONPG) was added in each well. The absorbance at 420 nm was measured using microtiter plate reader for 30 minutes to determine the appearance of the degradation product of ONPG (Vmax). Specific activities were calculated by dividing the Vmax of the OD_420_ appearance by the OD_600_, and the resulting values were multiplied by 100000 to approximate Miller units (empirical determination).

### Electrophoretic mobility shift assay

RNAs were transcribed in vitro from PCR products using primers listed in Table 2 and the MEGAShortScript T7 kit (Thermo Fisher Scientific) according to the manufacturer’s recommendations. RNAs were gel-purified, eluted and precipitated with ethanol in the presence of 0.3 M sodium acetate. RNAs were dephosphorylated using CIP enzyme and 6 pmols were labelled at the 5′ ends with [ψ-^32^P] with PNK enzyme. RNAs were denatured in buffer (330 mM KCl, 80 mM HEPES pH 7.5, 4 mM MgCl_2_) for 5 min at 80 °C and then refolded gradually at 30 °C for 10 min. Binding was conducted in 10 µL for 30 min at 37 °C. 0.3 pmol of phospho-labelled *iscR* mRNA were incubated with increasing amounts of unlabeled OxyS or FnrS sRNA. Samples were supplemented with glycerol to a final concentration of 10 %, then loaded onto native 6 % polyacrylamide gels (29:1) containing 5 % of glycerol. Electrophoresis was performed in Tris–borate–EDTA buffer (TBE) 0.5 X at 4 °C during 2 hours at 80 V. Results were revealed using a PhosphorImager, Typhoon FLA 9500 scanner (GE Healthcare).

### Antibiotic sensitivity experiments

Starting from anaerobic overnight cultures in LB, strains were diluted 1/100 times in fresh medium and grown aerobically at 37 °C with shaking until OD_600_ :: 0.2. Then, antibiotics were added to the culture (gentamicin: 5 µg / mL; kanamycine: 7 µg / mL). Samples were taken after 90 min and 180 min, serially diluted in PBS buffer and spotted on LB agar plates with or without bovine liver catalase (2000 U/plate; Sigma-Aldrich) and incubated at 37 °C for 16 h. Bacterial survival was determined by colony forming unit (C.F.U.) counting after overnight growth. CFU at time 0 was of approximatively 2 x 10^7^ cells / mL in all experiments.

### Minimum inhibitory concentration (MIC) determination

The MIC was determined in accordance with previous protocols (39). In brief, 100 µL of bacterial culture containing 2 x 10^5^ CFU / mL was inoculated into each well of a 96-wells microtiter plate that contained varying concentrations of antibiotics (gentamicin, kanamycin, tetracycline, ampicillin and norfloxacin). The plate was incubated aerobically at 37 °C for 18 h. The absorbance of each well was then measured at OD_600_ on a microtiter plate reader. The lowest concentration of the drug that entirely suppressed microbial growth was defined as the MIC.

### RNA extraction and northern blot experiments

Overnight cultures of the appropriate strains were diluted in fresh medium containing ampicillin (100 µg/mL) when indicated and incubated at 37 °C. At OD_600_ = 0.6, 1 mL samples of the cultures were collected and IPTG (100 µM) was immediately added to the culture, then new samples were extracted at indicated time points. RNA was extracted using hot-phenol method as previously described (46) and resuspended in 10 µL of diethyl pyrocarbonate (DEPC)-treated water. Total RNAs were run on 1.75 % agarose denaturing gels and was then transferred onto Zeta Probe (Bio-Rad) positively charged membranes by an overnight capillary transfer. RNAs were cross-linked to the membrane using a UV cross-linker. Membranes were then hybridized with specific biotinylated probes overnight at 38 °C, and RNAs were detected using North2South (Thermo Scientific) labeling kit according to manufacturer’s instructions. Oligonucleotides used as probes are described in Table 2. A probe against the abundant SsrA RNA was used as a control.

### Fluorescence intensity determination

Fluorescence intensity determination of bacterial cultures was performed using a TECAN Spark microtiter plate reader. Briefly, after a 1/100 dilution of an overnight culture, cells were grown with agitation at 37 °C and OD_600_ and the Gfp (excitation at 485 nm, emission at 535 nm) and or mCherry (excitation at 580 nm and emission at 620 nm) fluorescence intensity were measured at regular intervals over time or after 5 hours of culture. The specific fluorescence intensity was calculated by subtracting the autofluorescence of a non-fluorescent strain to that of the strains containing fluorescent fusion. In the case of Hpx^-^ strains, bacteria were cultivated overnight anaerobically, diluted at 1/100 in fresh anaerobic medium and cultivated anaerobically until DO_600_ = 0.3. Then cultures were used to inoculates a microtiter plate and cells were grown aerobically, with agitation at 37 °C to induce oxidative stress. The OD_600_, GFP and/or mCherry fluorescence intensity were measured over time in a TECAN Spark reader. For the study of FnrS in anaerobiosis, aerobic overnights were diluted (1/100) in Hungate’s tube containing LB rich medium and grown for 16 h. Then, 5 mL of the aerobic culture were centrifugated at 6000 g for 10 minutes and the pellet resuspended in 500 µL of PBS and kept at RT for 30 minutes in order to allow proper GFP folding. Then OD_600_ and GFP fluorescence intensity were determined in microtiter plate using TECAN spark apparatus.

## Acknowledgments

We would like to thank Gisela Storz for providing strains and precious advices along the course of this study. We deeply thank Ciaran Condon, Delphine Allouche, Jonathan Jagodnik and Noella Amiot for help with EMSA experiments. We would like to thank also Samuel Carien for plasmid construction and Yohann Duverger for technical advices and for reading the manuscript.

This work was funded by the Centre National de la Recherche Scientifique (CNRS, http://www.cnrs.fr), Aix Marseille Université (AMU, https://www.univ-amu.fr) and by the Agence Nationale de la Recherche (ANR-21-CE12-002, Kinebiotics). CB received fundings from the Fondation pour la Recherche Médicale (number: ECO201906009080 and FDT202204014707).

