## Supplementary material for "Small RNA regulation of an essential process induces bacterial resistance to aminoglycosides during oxidative stress"

**Figure Supp Legends**

**Figure S1: Growth of the WT and ∆*oxyS* strains containing the $P_{isc}$-*iscR-gfp* fusion and overexpressing sRNAs.** Growth curves of the WT (CB47) **(A)** and ∆*oxyS* (CB176) **(B)** strains containing the $P_{isc}$-*iscR-gfp* fusion transformed with the pBR-plac vector (plac, grey), pOxyS (red) and pFnrS (green) cultivated in LB containing 100 µM IPTG were determined by following the optical density at 600 nm over time.

**Figure S2: GFP fluorescence intensity measurements in strains CB47 and CB176 in absence of IPTG. (A-B)** GFP fluorescence intensity overtime of strains CB47 (A) and CB176 (B) containing the pBR-plac vector (plac, grey), pOxyS (red) and pFnrS (green) plasmids in LB culture without IPTG addition. **(C-D)** Growth curve corresponding to experiments in panels A and B, respectively, determined by following the optical density at 600 nm over time.

**Figure S3: GFP fluorescence intensity measurements in the CB56 strain containing the psRNA. (A)** Cultures of the CB56 strain containing the P*isc-iscS-gfp* fusion transformed with the plasmids plac (grey), pOxyS (red), pFnrS (green) and pRyhB (blue) were performed in LB and GFP fluorescence intensity was followed overtime, in absence of IPTG. **(B-C)** Growth curves of the CB56 strains containing plac, pOxyS, pFnrS and pRyhB in absence (B) or in presence (C) of 100 µM IPTG. Growth was determined by following absorbance at $OD_{600}$ over time.

**Figure S4: OxyS base-pairs to the *iscR* mRNA. (A)** Native EMSA of $^{32}$P-labeled *iscR* full length (FL) (0.2 pmole) with increasing amounts of OxyS. The white arrow indicates the *iscR* FL RNA and the black arrow the complex formed between the *iscR* FL RNA and OxyS **(B)** Native EMSA of $^{32}$P-labeled *iscR* 5' (0.3 pmole) and increasing amount of WT OxyS RNA or OxyS mut (GCC -> AAA) RNA. The white arrow indicates free *iscR 5** and the black arrow indicates the *iscR 5** - OxyS complex.

**Figure S5: FnrS control of *iscR* expression in anaerobic conditions. (A).** Northern blots were performed from RNA extractions of overnight cultures of the PM1490 strain (WT) or its ∆*fnrS* derivative (PM3106) in LB medium in presence of absence of oxygen (Hungate's tubes). The membrane was hybridized with a probe against the FnrS RNA or SsrA as a control. **(B)** FnrS regulation of $P_{BAD}$-*iscR-lacZ* fusion expression. The effect of the deletion of *fnrS* (green) on *iscR'-'lacZ* fusion compared to the WT (grey). Both strains were grown anaerobically as in (A).

**Figure S6: Growth curves of the $Hpx^-$ strain containing the P*isc-iscR-gfp* fusion.** Growth curves of the $Hpx^-$ strain containing the P*isc-iscR-gfp* fusion (CB182, grey) and its isogenic ∆*oxyS* derivative (CB186, red) containing *iscR'-'gfp* fusion during oxidative stress. Strains were grown anaerobically (0 min) and then transferred aerobically leading to oxidative stress induction. Growth was determined by following the absorbance at $OD_{600}$ measured every 15 minutes during 16 hours after oxidative stress induction.

**Figure S7**: ***IscR* expression in absence of oxidative stress. (A-B)** The CB47 strain containing the $P_{isc}$-*iscR-gfp* (grey) and its Δ*oxyS* derivative (CB58, red) were cultured anaerobically and then transferred aerobically in the same way as with the $Hpx^-$ and $Hpx^-$ Δ*oxyS* strains in Figure S7. Fluorescence intensity (A) and growth ($OD_{600}$) (B) were measured every 15 minutes during 15 hours after the switch to aerobic conditions.

**Figure S8: Growth curves of $Hpx^-$ strain containing *sufA'-'mCherry* fusion in presence of oxidative stress.** Growth curves of $Hpx^-$ strain containing *sufA'-'mCherry* fusion (CB190, grey) and its Δ*oxyS* isogenic derivative (CB200, red) during oxidative stress. Strains were grown and growth was measured in the same way as in Fig. S7.

**Figure S9: Conservation of *iscR, iscU, sufB, oxyS, fnrS* and *ryhB* across enterobacteria.** Sequence conservation of *iscR* (pale blue), *iscU* (magenta), *sufB* (yellow), *oxyS* (red) and *ryhB* (orange) from *E. coli* MG1655 was performed using NCBI BLAST (https://www.ncbi.nlm.nih.gov/) against the RefSeq Representative genomes database, restricted to enterobacteria and using default parameters. Number in brackets next to the genera names represent the number of total representative genomes represented. Color intensity correspond to the presence of the genes in the different genomes; absence of color means that the gene is not present in the represented genera.

**Table S1: Strains and plasmids used in this study.**

| Strain | Description | Reference |
|---|---|---|
| **MG1655** | *Escherichia coli K-12 substr. MG1655* | Parental strain |
| **BEFB20** | MG1655 *nuo::npt1* Δ*sdhB* | 1 |
| **GSO35** | *Escherichia coli* K-12 Δ*oxyS2::cm* | 2 |
| **Hpx⁻** | MG1655 Δ*ahpCF* Δ*katE* Δ*katG* | 3 |
| **PM1205** | MG1655 *lacI*::$P_{BAD}$:*cat-sacB*:*lacZ*, Δ*araBAD*, *araC*+, *mal::lacI^q*, mini $\lambda^{tet}$ | 4 |
| **PM2250** | MG1655 *mal::lacI^q*, *lacI'*::$P_{iscR}$-*iscR'-'lacZ* | This study |
| **PM3030** | MG1655 *mal::lacI^q*, Δ*araBAD*, *lacI'*::$P_{BAD}$-*iscR'-'lacZ* | This study |
| **CB8** | MG1655 *oxyS::cat* | This study |
| **PM3106** | MG1655 *fnrS*::*tet* | This study |
| **GSO402** | MG1655 *fnrS::kan* | 5 |
| **CB47** | MG1655 *mal::lacI^q*, *lacI'*::$P_{isc}$-*iscR'-'gfp* | This study |
| **CB58** | CB47 *oxyS::cat* | This study |
| **CB56** | MG1655 *mal::lacI^q*, *iscS'-'gfp-kan* | This study |
| **CB176** | CB47 *oxyS::cat*, *fnrS::kan*, *ryhB::spc* | This study |
| **CB178** | Hpx⁻ *oxyS::cat* | This study |
| **CB182** | Hpx⁻ *lacI'*::$P_{isc}$-*iscR'-'gfp-kan* | This study |
| **CB186** | CB182 *oxyS::cat* | This study |
| **CB203** | CB182 *oxyS::cat*, *fnrS::tet*, *ryhB::spc* | This study |
| **CB190** | Hpx⁻ $P_{suf}$-*sufA-mcherry-kan* | This study |
| **CB200** | CB190 *oxyS::cat* | This study |
| **CB204** | CB182 *mal::lacI^q*, *oxyS::cat*, *ryhB::spc*, *fnrS::tet* | This study |
| **CB226** | Hpx⁻ *iscR::kan* | This study |
| **CB228** | Hpx⁻ *oxyS::cat*, *iscR::kan* | This study |
| **CB390** | Hpx⁻ *nuo*::*npt1* | This study |
| **CB394** | Hpx⁻ *oxyS::cat*, *nuo::npt1* | This study |
| **CB433** | PM3030 *oxyS::cat, fnrS::kan, ryhB::spc* | This study |

| Plasmid | Description | Reference |
|---|---|---|
| **pBR-plac** | pBR22 derivative carrying a modified $P_{lacO-1}$ promoter, $Amp^R$ | 6 |
| **pOxyS** | $Amp^R$, Aat-II-EcoR1 *oxyS* cloned in pBR-plac | 4 |
| **pFnrS** | $Amp^R$, Aat-II-EcoR1 *fnrS* cloned in pBR-plac | 4 |
| **pRyhB** | $Amp^R$, Aat-II-EcoR1 *ryhB* cloned in pBR-plac | 4 |
| **pOxyS mut** | $Amp^R$, Aat-II-EcoR1 *oxyS* (39-41 GCC>AAA) in pBR-plac; synthetized by Azenta | This study |
| **pTrc99a** | *lacI^q*, pUC18 EcoRI-HindIII polylinker, $Amp^R$ | 7 |
| **piscR** | $Amp^R$, *Nco*I-*Bam*HI *iscR* cloned in pTrc | This study |
| **pGBMKN-GFP** | $Spc/Sm^R$ | 8 |
| **pGBMKn-mCherry** | $Spc/Sm^R$ | 8 |
| **pkD46** | oriR101, repA101ts, *araC*, $P_{araB}$ , $Ap^R$; | 9 |
| **pCP20** | $Amp^R$, $Cat^R$, cI857, $ori^{Ts}$, λPR, *flp* | 10 |


1. Ezraty, B. *et al.* Fe-S cluster biosynthesis controls uptake of aminoglycosides in a ROS-less death pathway. *Science* **340**, 1583–1587 (2013).
2. Altuvia, S., Weinstein-Fischer, D., Zhang, A., Postow, L. & Storz, G. A small, stable RNA induced by oxidative stress: role as a pleiotropic regulator and antimutator. *Cell* **90**, 43–53 (1997).
3. Ezraty, B., Henry, C., Hérisse, M., Denamur, E. & Barras, F. Commercial Lysogeny Broth

culture media and oxidative stress: a cautious tale. *Free Radic Biol Med* **74**, 245–251 (2014).
4. Mandin, P. & Gottesman, S. A genetic approach for finding small RNAs regulators of genes of interest identifies RybC as regulating the DpiA / DpiB two component system. *Mol Microbiol* **72**, 551–565 (2009).
5. Durand, S. & Storz, G. Reprogramming of anaerobic metabolism by the FnrS small RNA. *Mol. Microbiol.* **75**, 1215–1231 (2010).
6. Guillier, M. & Gottesman, S. Remodelling of the Escherichia coli outer membrane by two small regulatory RNAs. *Mol Microbiol* **59**, 231–247 (2006).
7. Amann, E., Ochs, B. & Abel, K. J. Tightly regulated tac promoter vectors useful for the expression of unfused and fused proteins in Escherichia coli. *Gene* **69**, 301–315 (1988).
8. Mercier, R. *et al.* The MatP/matS site-specific system organizes the terminus region of the E. coli chromosome into a macrodomain. *Cell* **135**, 475–485 (2008).
9. Datsenko, K. A. & Wanner, B. L. One-step inactivation of chromosomal genes in Escherichia coli K-12 using PCR products. *Proceedings of the National Academy of Sciences* **97**, 6640–6645 (2000).
10. Cherepanov, P. P. & Wackernagel, W. Gene disruption in Escherichia coli: TcR and KmR cassettes with the option of Flp-catalyzed excision of the antibiotic-resistance determinant. *Gene* **158**, 9–14 (1995).

**Table S2: Oligonucleotides used in this study**

| Name | Sequence (5’ – 3’) |
|---|---|
| **iscR-GFP-F** | GAAGTAAGACATGAGACTGACATCTAAAGGGCGCTATGCCGAATCTGTGAGCAAGGGCGAGGAGCT |
| **lacZ-GFP-Kan-R** | TTACGCGAAATACGGGCAGACATGGCCTGCCCGGTTATTAGTCCATATGAATATCCTCCTTAG |
| **PBAD-iscR-F** | ACCTGACGCTTTTTATCGCAACTCTCTACTGTTTCTCCATGCTATGCAATACCCCCACTT |
| **lacZ-iscR-R** | TAACGCCAGGGTTTTCCCAGTCACGACGTTGTAAAACGACCACGGCATAGCGCCCTTTAG |
| **Pisc-F** | CGAAGCGGCATGCATTTACGTTGACACCATCGAATGGCGCTACAGTGAACAGAACCGTAG |
| **iscS-GFP-F** | GCAGGGCGTGGATCTGAACAGCATCGAATGGGCTCATCATGAATCTGTGAGCAAGGGCGAGGAGCT |
| **iscS-GFP-R** | CGCTGTAAGCCATTATAAATTCTCCTGATTCCGATACCGAGTCCATATGAATATCCTCCTTAG |
| **sufA-mcherry-F** | AGCCCAGAATGAATGTGGCTGTGGCGAAAGCTTTGGGGTAGAATCTGTGAGCAAGGGCGAGGAG |
| **sufA-mcherry-R** | ACATCGTCAGTTGCTTCAGTATTACGAGACATAGTACCGCGAACTCCAGCATGAGATCCCCGCGC |
| **iscR-T7-F** | GCTCTAATACGACTCACTATAGACAATAAAAAACCCCGGGCAGGGGCGAGTTTG |
| **iscR-T7-R** | GGAAATATCAGCCAACGGTACC |
| **iscR-5’-T7-F** | GCTCTAATACGACTCACTATAGGCTATGCAATACCCCCACTTTTAC |
| **iscR-5’-RT** | GATGTCAGTCTCATGTCTTAC |
| **iscRdown-BamHI** | CGCGGATCCTTAAGCGCGTAACTTAACGTC |
| **NcoI-iscRup** | GCCCCATGGCAAGACTGACATCTAAAGGGCGC |
| **SsrA NB** | CGCCACTAACAAACTAGCCTGATTAAGTTTTAACGCTTCA |
| **OxyS NB** | AAACTCTCGAAACGGGCAGTGACTTCAAGGGTTAAA |
| **FnrS NB** | GGAAGTAAGACAATATGGAGCGCAACGCCCATCGC |
| **RyhB NB** | AAGTAATACTGGAAGCAATGTGAGCAATGTCGTGCTTTCAGGTTCTC |
| **iscR NB** | CGCAACGTCAAGCATTGCGGTCACGGCATAGCGCCCTTTAGATGTCAGTC |

### Figure S1

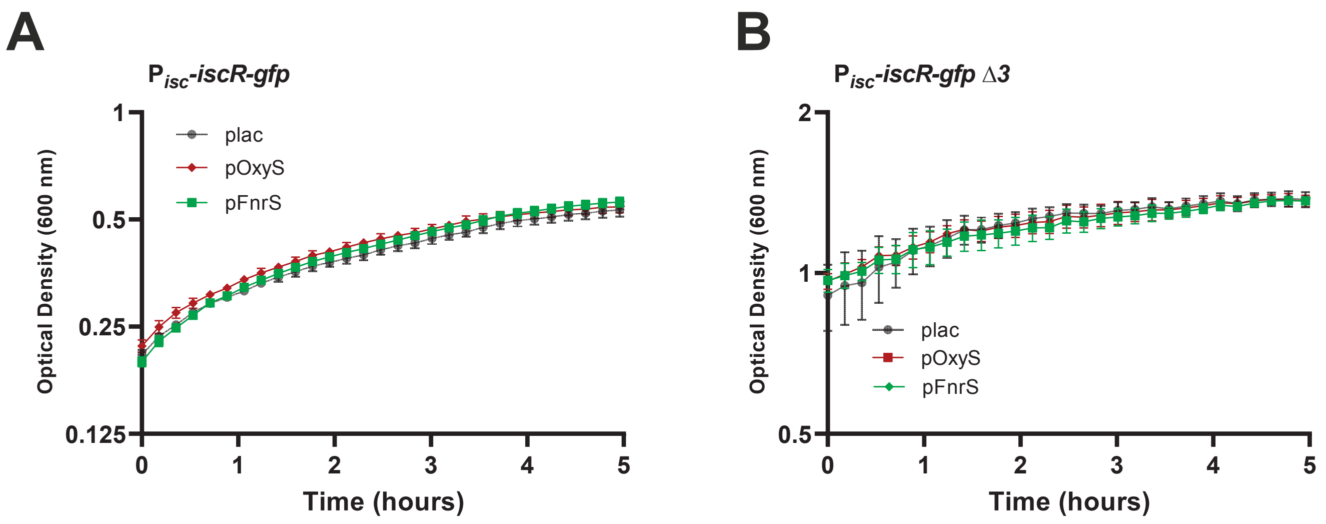

A
$P_{isc}$-*iscR-gfp*
Optical Density (600 nm)
1
0.5
0.25
0.125
plac
pOxyS
pFnrS
0
1
2
3
4
5
Time (hours)
B
$P_{isc}$-*iscR-gfp* Δ3
Optical Density (600 nm)
2
1
0.5
plac
pOxyS
pFnrS
0
1
2
3
4
5
Time (hours)

### Figure S2

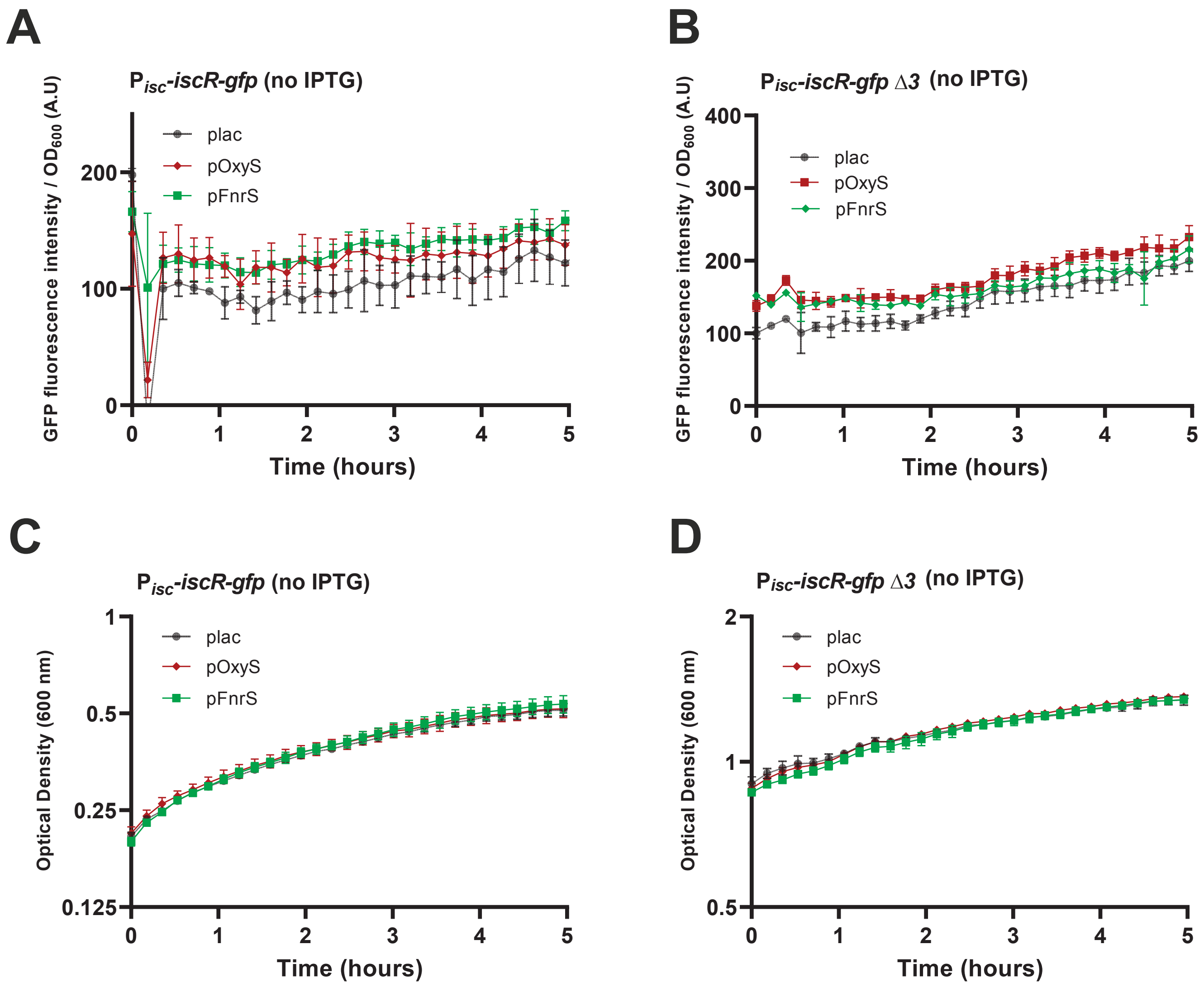
A
$P_{isc}$-iscR-gfp (no IPTG)
GFP fluorescence intensity / $OD_{600}$ (A.U)
Time (hours)
plac
pOxyS
pFnrS
B
$P_{isc}$-iscR-gfp Δ3 (no IPTG)
GFP fluorescence intensity / $OD_{600}$ (A.U)
Time (hours)
plac
pOxyS
pFnrS
C
$P_{isc}$-iscR-gfp (no IPTG)
Optical Density (600 nm)
Time (hours)
plac
pOxyS
pFnrS
D
$P_{isc}$-iscR-gfp Δ3 (no IPTG)
Optical Density (600 nm)
Time (hours)
plac
pOxyS
pFnrS

### Figure S3

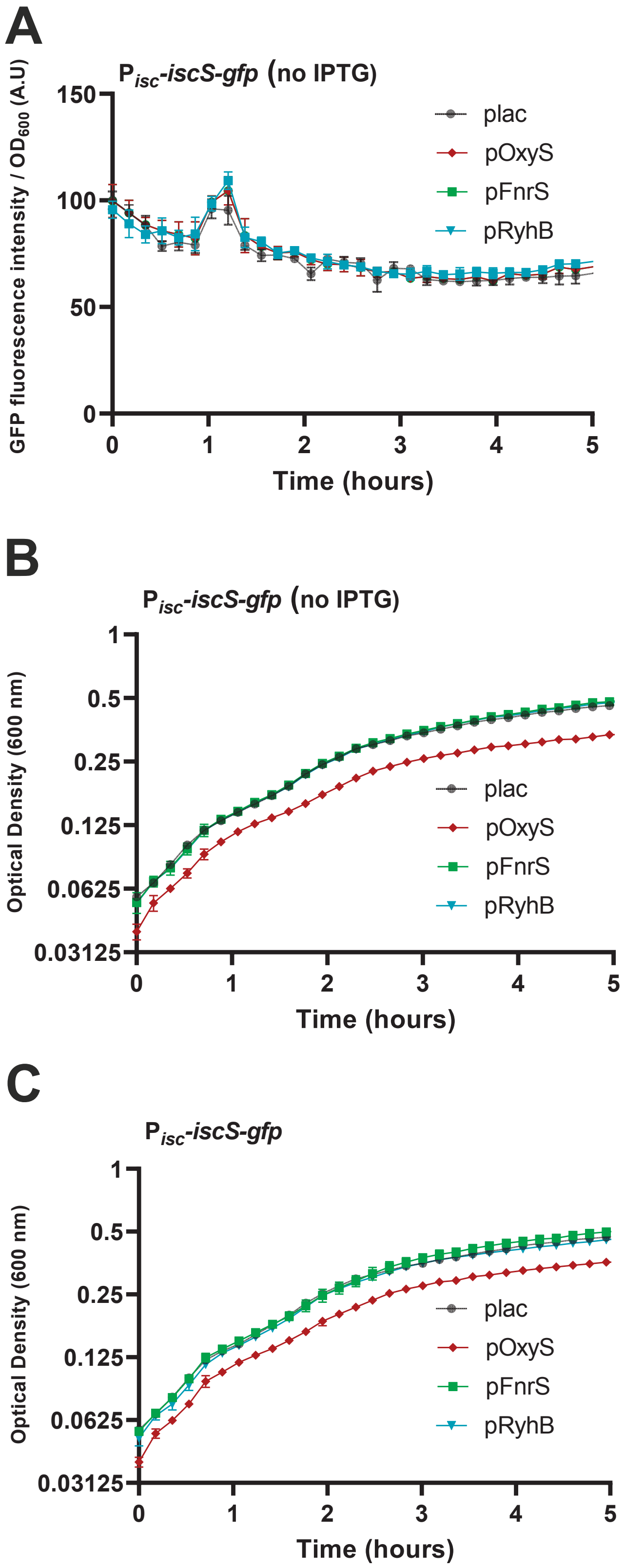

A
$P_{isc}$-*iscS-gfp* (no IPTG)
GFP fluorescence intensity / $OD_{600}$ (A.U)
150
100
50
0
0
1
2
3
4
5
Time (hours)
plac
pOxyS
pFnrS
pRyhB
B
$P_{isc}$-*iscS-gfp* (no IPTG)
Optical Density (600 nm)
1
0.5
0.25
0.125
0.0625
0.03125
0
1
2
3
4
5
Time (hours)
plac
pOxyS
pFnrS
pRyhB
C
$P_{isc}$-*iscS-gfp*
Optical Density (600 nm)
1
0.5
0.25
0.125
0.0625
0.03125
0
1
2
3
4
5
Time (hours)
plac
pOxyS
pFnrS
pRyhB

### Figure S4

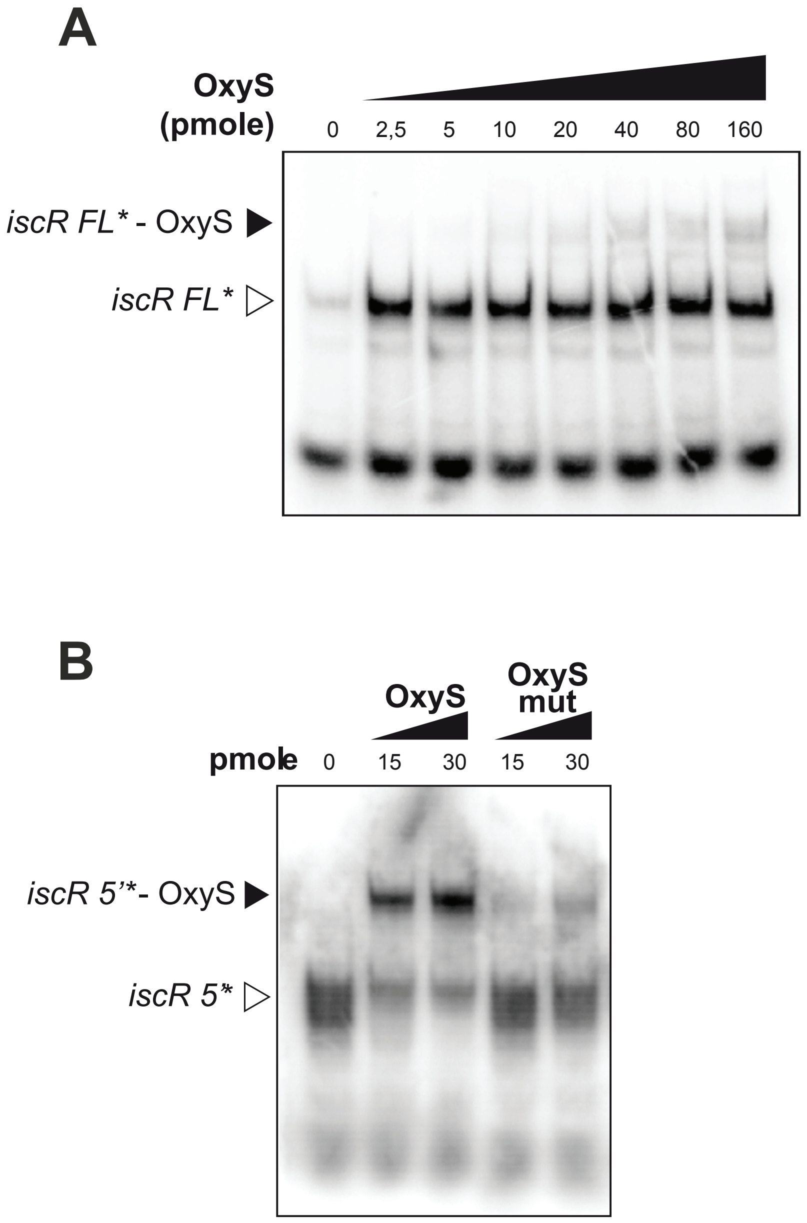

A
OxyS
(pmole)
0
2,5
5
10
20
40
80
160
iscR FL* - OxyS
iscR FL*
B
OxyS
OxyS
mut
pmole
0
15
30
15
30
iscR 5'*- OxyS
iscR 5*

### Figure S5

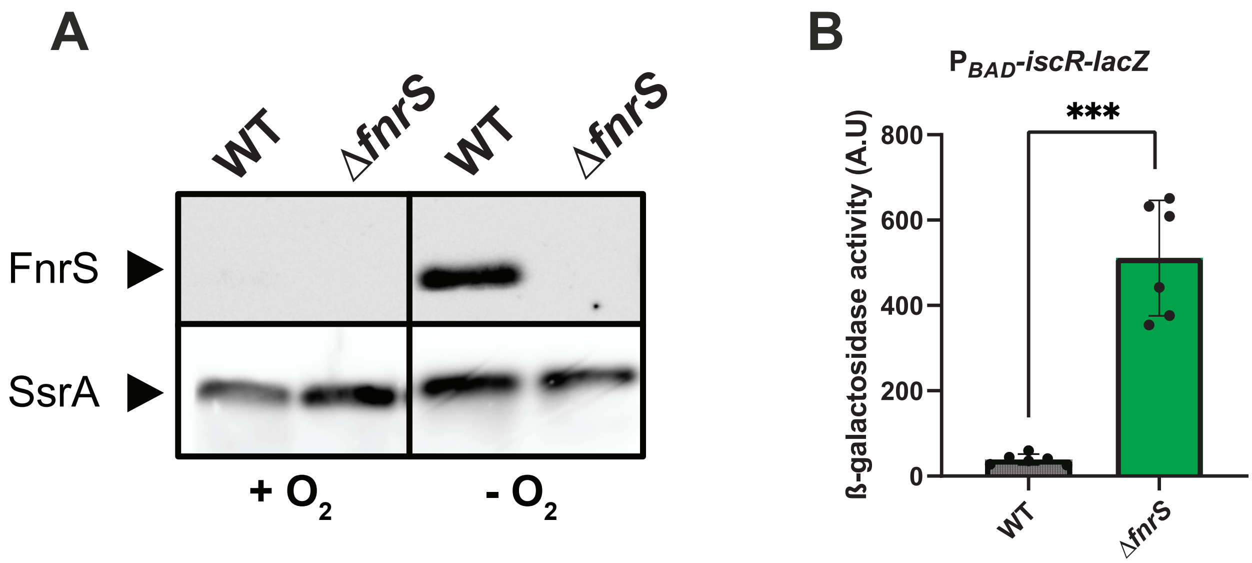
A
WT
ΔfnrS
WT
ΔfnrS
FnrS
SsrA
+ O2
- O2
B
PBAD-iscR-lacZ
***
ß-galactosidase activity (A.U)
800
600
400
200
0
WT
ΔfnrS

### Figure S6

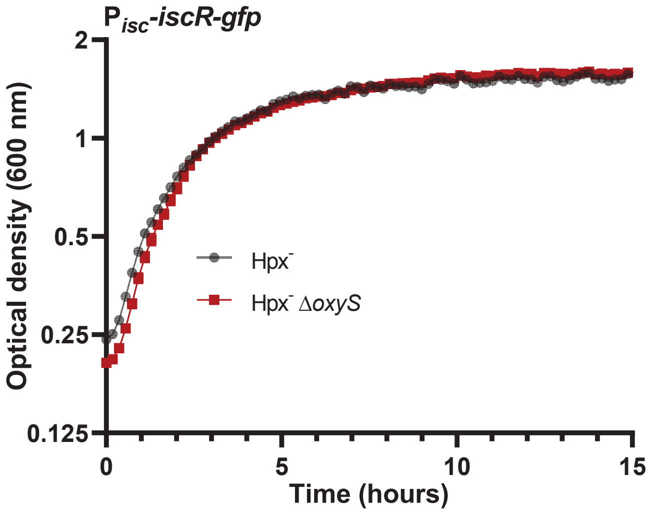

P_isc-iscR-gfp
Optical density (600 nm)
2
1
0.5
0.25
0.125
0
5
10
15
Time (hours)
Hpx-
Hpx- ΔoxyS

### Figure S7

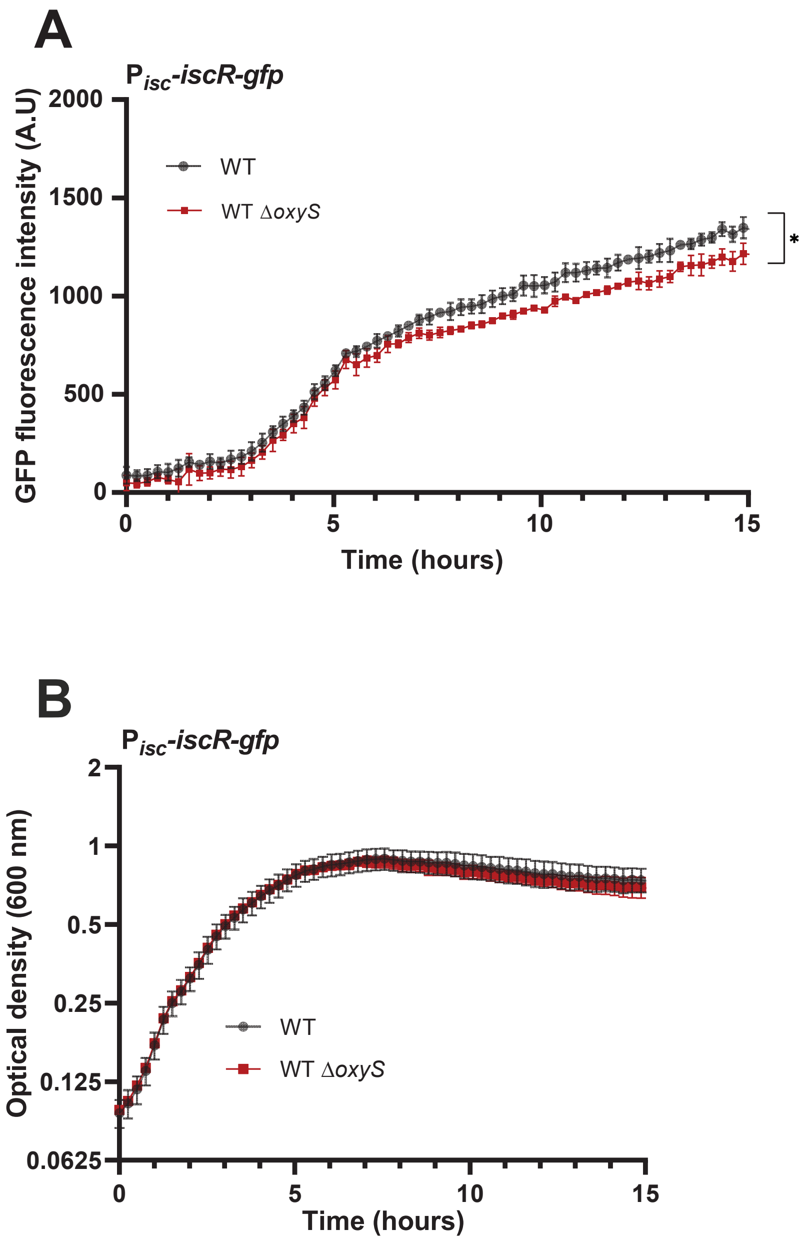
A
$P_{isc}$-iscR-gfp
GFP fluorescence intensity (A.U)
2000
1500
1000
500
0
WT
WT ΔoxyS
*
0
5
10
15
Time (hours)
B
$P_{isc}$-iscR-gfp
Optical density (600 nm)
2
1
0.5
0.25
0.125
0.0625
WT
WT ΔoxyS
0
5
10
15
Time (hours)

### Figure S8

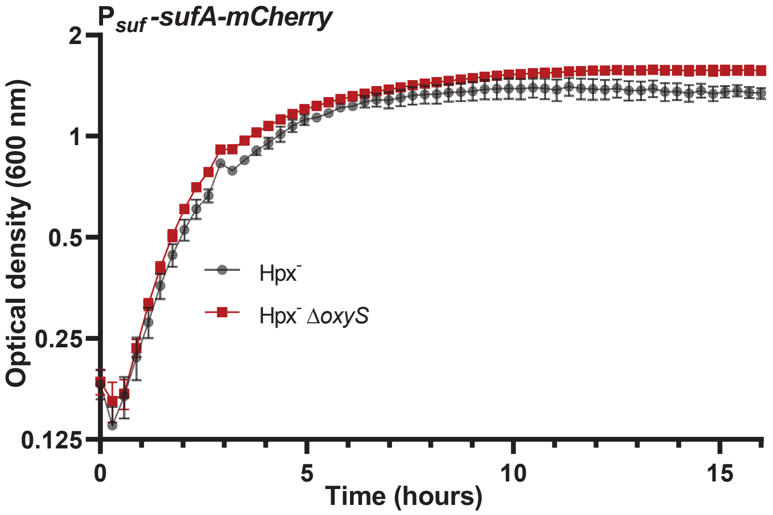

P suf -sufA-mCherry
Optical density (600 nm)
2
1
0.5
0.25
0.125
0
5
10
15
Time (hours)
Hpx-
Hpx- ΔoxyS

**Figure S9**

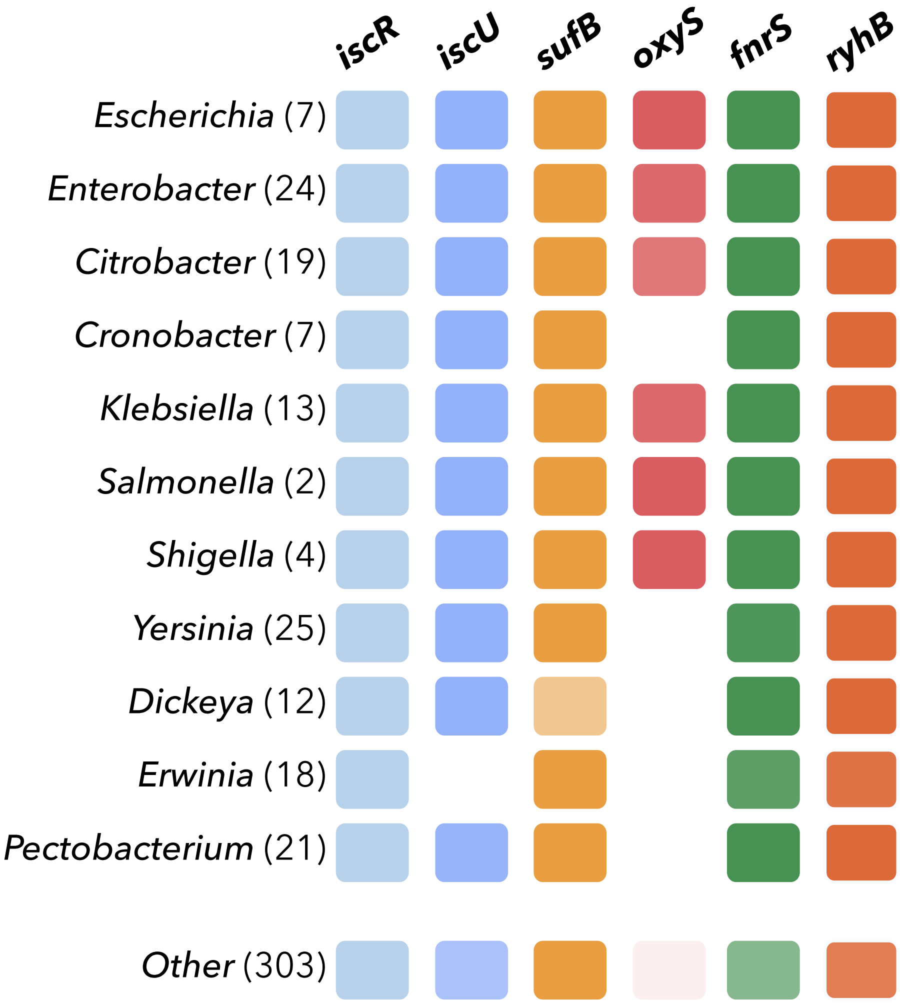
iscR
iscU
sufB
oxyS
fnrS
ryhB
*Escherichia* (7)
*Enterobacter* (24)
*Citrobacter* (19)
*Cronobacter* (7)
*Klebsiella* (13)
*Salmonella* (2)
*Shigella* (4)
*Yersinia* (25)
*Dickeya* (12)
*Erwinia* (18)
*Pectobacterium* (21)
*Other* (303)
